# Multiscale mechanical model for cell division orientation in developing biological systems

**DOI:** 10.1101/785337

**Authors:** B. Leggio, J. Laussu, E. Faure, P. Lemaire, C. Godin

**Author notes:** Institut Clément Ader, Université Toulouse III, CNRS, INSA, ISAE-SUPAERO, Mines-Albi, France.

## Abstract

Developing biological structures are highly complex systems, within which shape dynamics at different places is tightly coordinated. One key process at play during development is the regulation of cell division orientation. In this work, through a reformulation of cell division in terms of its energetic cost, we propose that oriented cell division is one mechanism by which cells can read and react to mechanical forces propagating in a tissue even in the absence of interphase cellular elongation in the direction of these forces. This view reproduces standard geometric division long-axis rules as a special case of a more general behaviour, in which systematic deviations from these rules can emerge. We show that states of anisotropic tension in multicellular systems can be the cause of these deviations, as often experimentally found in living tissues. Our results provide a unifying view on the different intracellular mechanisms at play in orienting cell division: they are processes which minimize energy loss, reflecting a trade-off between local and long-range mechanical signals. The consequences of this competition are explored in simulated tissues and confirmed *in vivo* during both the development of the pupal epithelium of dorsal thorax in *D.* melanogaster and the epidermal morphogenesis of ascidian embryos.

**Author summary:** In this work we reformulate the process of cell division orientation in development as a mechanical-energy optimization. We show that classical rules for division orientation naturally emerge when a cell minimizes the work performed against its local environment. Moreover, when multicellular stress profiles are taken into account, observed systematic violations of these rules are explained in correlation with states of anisotropic tension within the tissue. We confirm our findings experimentally on developing systems imaged with cellular resolution. Our results provide a new paradigm to understand cell division in multicellular contexts and contribute to building a physical view of biological phenomena.

## I. INTRODUCTION

During animal development, morphogenesis is driven by dynamical processes occurring at specific positions.

These local processes, however, change the global shape of the system, indicating the presence of mechanisms through which local shape changes are transmitted to - and interpreted by - cells located at a distance [1–3]. In spite of the many works investigating the dynamics of shape change in living systems [4–12], and several studies investigating the nature of the information regulating these changes [13, 14], how such information propagates and is interpreted remains elusive.

Cell division orientation is one important component of morphogenesis. In animals it contributes for instance in shaping and elongating tissues, maintaining tissue homeostasis, generating cellular heterogeneity [15–19]. Cell division orientation is often correlated to cell shape. In both animals and plants, the orientation of a cell’s division plane (i.e., the direction orthogonal to the plane) often aligns with to the long geometrical axis of the cell at interphase-a rule known as Hertwig’s rule (HR) in animals [15, 20, 21] and Errera’s rule in plants [22, 23].

The division plane orientation is provided by the position of the mitotic spindle [24]. In animal cells, spindles are anchored to the cell cortex through astral microtubules emerging from the centrosomes and exerting mechanical forces on them. As a consequence, the spindle equilibrium position due to these forces has been shown to depend on cell geometry, which provides a physical explanation of long-axis rules for cell division [25–27]. It has further been shown that, in certain species [21, 28], microtubles anchoring at the cortex is biased by protein accumulation at tricellular junctions. This explains why, for instance, division orientation in the *Drosophila* epidermis is better predicted by the interphase distribution of tricellular junctions than by the cell shape.

More generally, the action of intra- or extra-cellular mechanical forces on cell division orientation has been extensively studied [12, 21, 23, 25–31]. In these studies, the predicted division orientation is in very good agreement with the direction of the cell’s long-axis. These results suggest that it is not directly cell shape, but rather complex biochemical and biomechanical processes which may be affected by cell shape, that orient cell division. Despite the generality of such laws, systematic and reproducible deviations from them are commonly observed experimentally and found to correlate with states of anisotropic tension in the surrounding tissue [32–35]. These deviating cells often coexist with cells following HR. To explain at once these two cell populations (respecting HR and deviating from it), the existence of alternative intracellular mechanisms for spindle positioning has been proposed, which can be biased by anisotropic tension in the tissue [6, 25, 33, 35].

In spite of these efforts, a unifying understanding of the regulation of division orientation is still largely missing.

In animals, cell division itself is often accompanied by radical active changes of shape as the cell proceeds to, through and out of mitosis [16, 36–39]. Mitosis usually does not affect the cell volume (although exceptions have been reported [36, 39]), but can lead to an increase in apical surface and in general alters cell form [38–41]. Frequently, an initial increase in cortical tension results in so-called mitotic rounding, which is then followed by cell elongation along the direction orthogonal to the division plane during the stage known as anaphase and, later, during the completion of cytokinesis. Thanks to the burst of new imaging technologies [42–45], the dynamics of cell shape during mitosis and its robustness against environmental and intrinsic noise can be now investigated in a quantitative manner.

In tightly-packed tissues, dividing animal cells push against their neighbours to create space to accommodate shape changes and volume increase during and after mitosis [36, 39]. If a cell’s environment is too stiff for the cell to deform it, the so-called spindle assembly checkpoint prevents progression to anaphase and stalls the division process [36]. Division is thus a process in which the cell’s surrounding feeds back information, used by the cell to control its mitosis. This implies that the dividing cell spends a certain amount of energy in changing its shape against the mechanical resistance of surrounding tissues. In [32], a mechanical model for the influence of direct neighbours on cell division orientation was given: neighbours can change a cell’s long axis due to mechanical forces depending on their topology. However, this model only accounted for the mechanical influence of a cell’s long-axis, assuming HR as the standard division rule. Also, no connection to multicellular mechanical stress patterns was considered. Yet, in light of the results of [36], the ability of a cell to deform its environment is a key information regulating the progression of its mitotic process. Since such deformation implies a flow of energy from the cell to its local neighbourhood under the form of mechanical work, we make the hypothesis that a strong connection exists between the regulation of cell division and the amount of a cell’s internal energy flowing to its environment through mechanical forces. However, no physical model has been developed yet to study this phenomenon: filling this gap is the main contribution of our work.

In this work, we developed a theoretical model of the energetic cost of cell division and its effect on cell division orientation. Our results show that cell division proceeds along the path that minimizes (mechanical) energy dissipation due to a cell’s interaction with its local and remote environments. This interaction provides the cell with information on the mechanical state of its surroundings. Such information is mechanically supplied to (and integrated by) a dividing cell and used as a cue to orient its division plane, which contributes to ensuring a global long-range coherence to evolving biological forms. Through this energetic view, we can explain at once HR and deviations from it as limiting cases of a general behaviour. We show that the competing intracellular mechanisms for division orientation suggested to explain deviations from HR, can be given a unifying view as optimal responses to minimize energy loss of the dividing cell. Hence, both the intracellular mechanisms responsible for HR and those biased by tissue tension allow for cellular energy optimisation.

## II. CELL DIVISION IN MULTICELLULAR SYSTEMS: AN ENERGETIC VIEWPOINT

Inspired by the shape dynamics observed during cell division in ascidian embryonic epidermis (Section IV of Supporting Information) and by the results of [36], we assumed here that the portions of space occupied by the mother cell just before mitosis and by the union of the two daughters just after mitosis differ. In ascidian embryonic epidermis, for instance, cell division is accompanied by an increase in the apical surface of the cellular clone (Fig. 4 of Supporting Information).

**FIG. 1:**
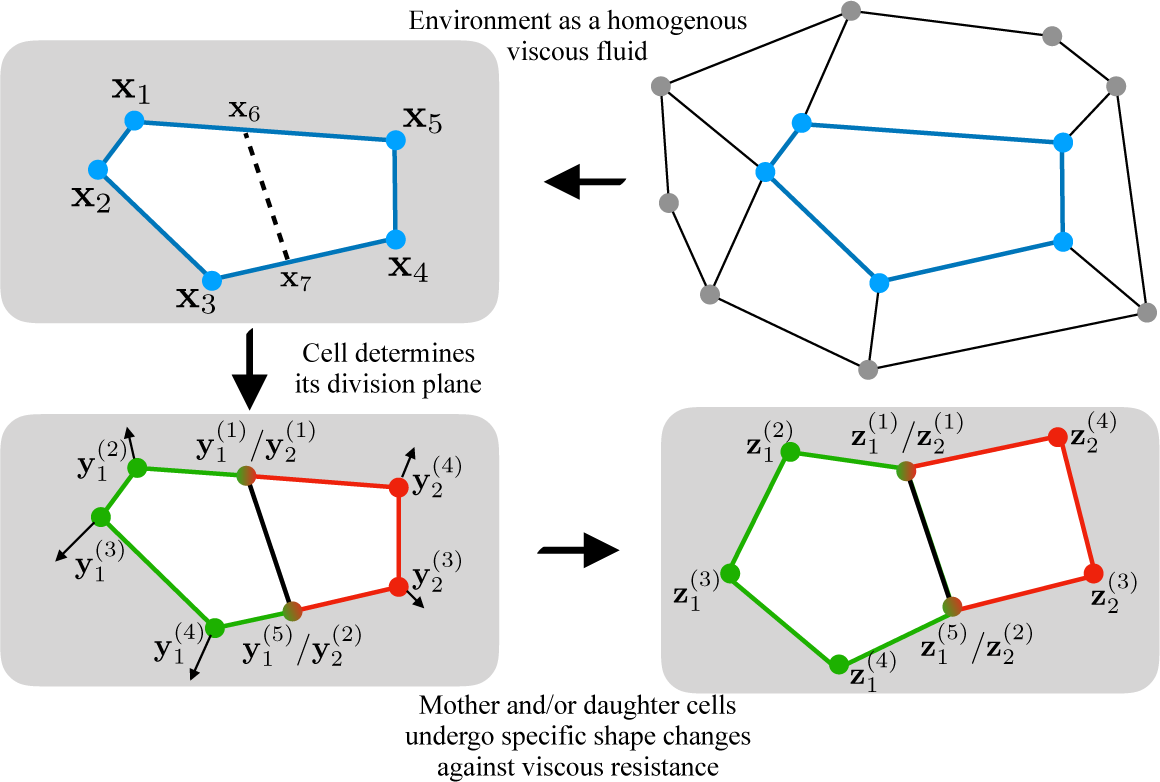
2D representation of cell division in a viscous environment. A cell with 5 neighbours and a given rest shape (vertex positions {**x**_*i*_}) divides, creating two daughter cells with, respectively, 5 and 4 neighbours and shapes given by vertices 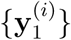 and 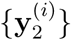. These two cells deform their surrounding environment to achieve specific regular shapes (for instance, spherical cells), given by vertex positions 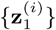 and 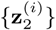. The latter depend on the orientation **n** of the division plane. Each vertex follows a trajectory 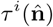 from its initial position **y**^(*i*)^ to its final position **z**^(*i*)^.

**FIG. 2:**
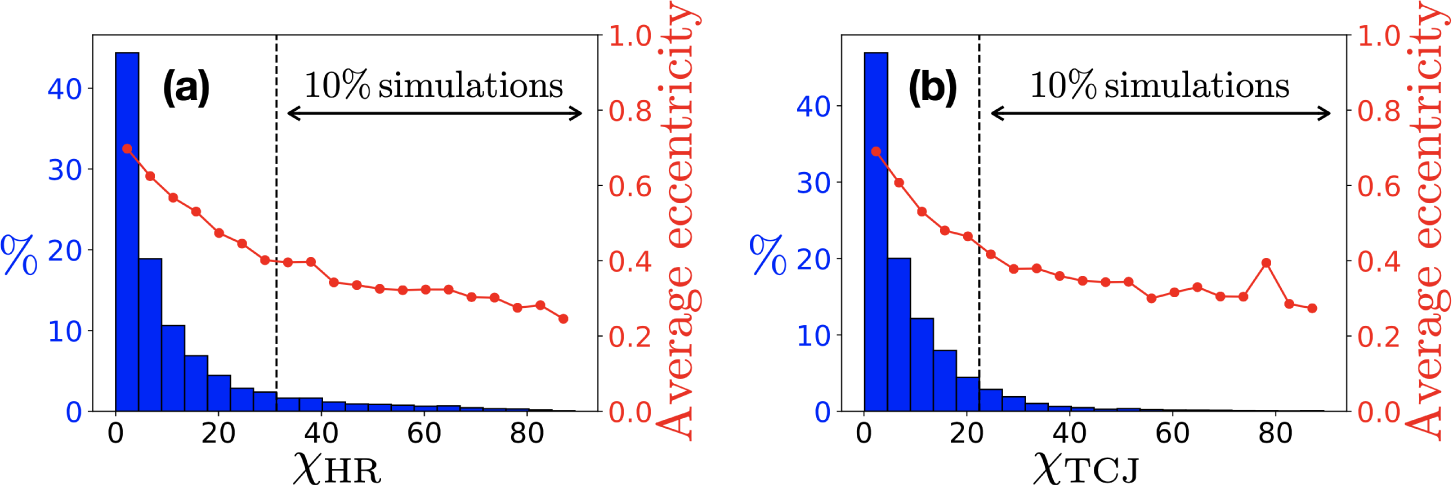
Distribution of deviation angles between predictions of Eq. (1) and cell main elongation ((a), 𝒳_HR_) or main direction of tricellular junctions distribution ((b), 𝒳_TCJ_). Specifically, 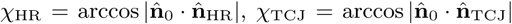. Results are obtained for 10^4^ randomly simulated cells. Here 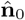 minimizes Eq. (1); 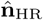 is the cell’s main elongation axis and 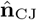 the main direction of its tricellular junctions. Red points (right vertical scale) give the average eccentricity of the ellipses approximating each cell in each histogram bar. 90% of cells have 𝒳_TCJ_ < 22° and 𝒳_HR_ < 30°, in agreement with what is found *in vivo* [21].

**FIG. 3:**
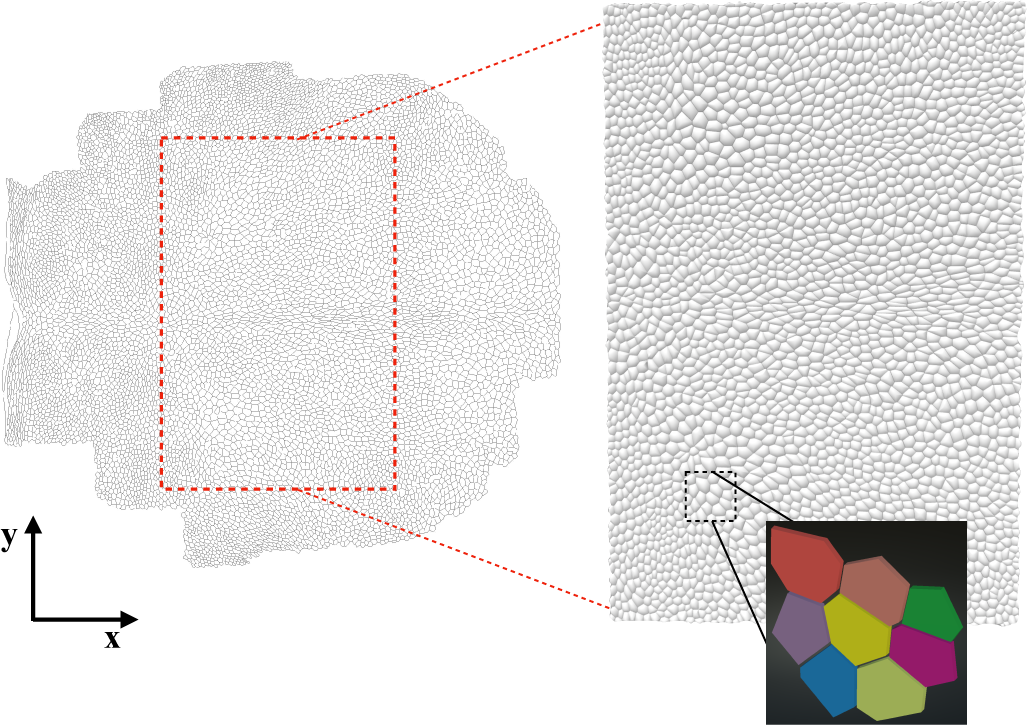
Reconstruction and polygonal approximation of the *Drosophila* pupal dorsal thorax epithelium. Data from [7].

**FIG. 4:**
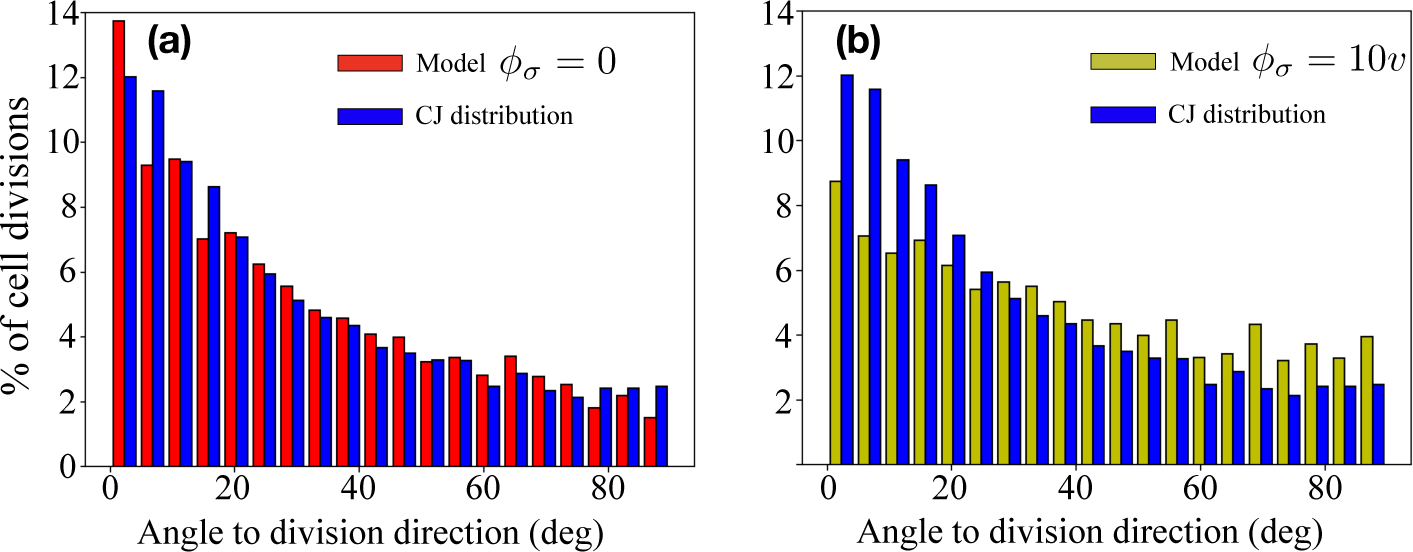
Distribution of deviation angles (deg) between TCJ directions and cell division orientation (blue bars), the predictions of Eq. (3) with *ϕ*_*σ*_ = 0 (red bars, panel (a)) and the predictions of Eq. (3) with *ϕ*_*σ*_ = 10*v* (yellow bars, panel (b)).

In our model of the energetics of cell division, we neglected the contribution of cell rounding around mitosis. The reason for this is that mitotic rounding of apical shape is not always the dominant shape-changing process: ascidian epidermal cells, as shown in Section IVB of the Supporting Information, show little mitotic rounding at the level of their apical surface compared to the subsequent surface elongation. While cells round up in 3D, their apical surfaces are only marginally affected by the process and tend to keep memory of their interphase shape until anaphase.

In general, the less important are shape changes during mitotic rounding-up compared to the ones at anaphase and cytokinesis, the more precise our formulation is.

We employed here a classical vertex-based representation of cellular shape [46], according to which the shape of a 2D cell is abstracted as the positions **R**_*i*_ of its junctions, represented as vertices of a polygon.

### A. The role of local cellular environment

Biological tissues are usually modelled as non-equilibrium viscoelastic mechanical systems [47], with properties changing with time and observation scale. In many circumstances one can, however, model the resistance offered by biological matter to displacement as a viscous fluid with viscous friction parameter *η*. This simplification allows to explicitly model the energetic cost of shape changes. Such changes happen against the viscosity of the surroundings, and as such must entail an energetic price paid by the internal energy of the cell. Interestingly, this view implies that the direction and amount of deformation needed depends on the orientation of the division plane with respect to the distribution of cell vertices.

Consider a dividing cell, described by its initial collection of vertices {**x**_*i*_} as represented in Fig. 1. Once mitotic spindles are positioned and the division plane is fixed, the cell is virtually composed of its two future daughters 1, 2 with vertices, respectively, 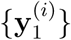 and 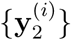. During and after division, cells change their shape by moving their vertices from initial positions 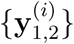 to final positions 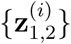, determined by the regular shape of the daughter cells (see Fig. 1). Energy is consumed by the viscosity of the medium due to viscous friction forces −*η***v** acting on each vertex moving with velocity **v**. We stress here that, due to our representation of cells as a collection of vertices, these viscous forces act on point particles (the vertices) rather than on extended objects. The work *W*_*η*_ of friction forces is

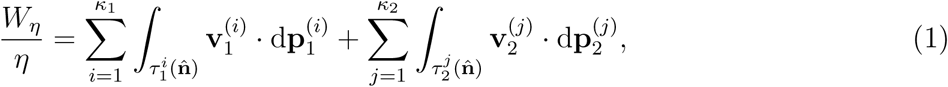

where 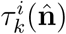, *k* = 1, 2 is the trajectory followed by the *i*-th vertex of daughter cell 1 (2), which depends on the division orientation 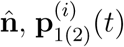 its instantaneous position along the trajectory, 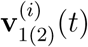 its finite velocity, 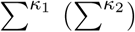 runs over all *κ*_1_ (*κ*_2_) vertices of daughter cell 1 (2) (common vertices contribute with weight 1*/*2 in both sums). Here and in what follows, a cell division orientation is defined as the direction orthogonal to the cell division plane (i.e., the direction orthogonal to the segment **x**_6_ − **x**_7_ in Fig. 1). The work in Eq. (1) depends on the amount of deformation required to accommodate the daughters’ final shapes. A natural consequence of this is that different division orientations correspond to different amounts of energy consumed by the dividing cell 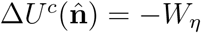. This formulation of the energetic cost of cell division (and its generalization, see Eq. (3) in the following) is the first contribution of this work.

### B. Long-range mechanical contribution to the energetics of cell division

Cells or tissues also actively change their shape, as a result of intra-cellular processes and/or the effect of external interactions [1, 2, 30, 47–50]. Like any movement of matter, an active local change in shape implies the build-up of mechanical forces within the tissue, whose distance of propagation depends on the properties of the cellular medium [47]. These forces create mechanical stress within the tissue, whose magnitude decreases with the distance from the center of the active process. We formalize these ideas considering a generic morphodynamic process *σ* at position **r**_*σ*_ (for instance, a cell actively changing its shape) which exerts a force **F**_*σ*_(**r, r**_*σ*_) = *ψ*_*σ*_(**r, r**_*σ*_)**ŵ**_*σ*_(**r, r**_*σ*_) at any position **r** in the tissue, of intensity *ψ*_*σ*_(**r, r**_*σ*_) and direction **ŵ**_*σ*_(**r, r**_*σ*_). Writing *ψ*_*σ*_ = *F*_*σ*_*h*(*d*(**r, r**_*σ*_)), where *F*_*σ*_ is the force intensity at the position of the process *σ, h*(*x*) is a monotonically decreasing function with *h*(0) = 1 (accounting for the decrease of stress magnitude with distance, due to the viscoplastic properties of living tissues [47, 51, 52]), *d*(**r, r**_*σ*_) is the euclidean distance between **r** and the process *σ* and **f**_*σ*_(**r, r**_*σ*_) = **ŵ**_*σ*_(**r, r**_*σ*_)*h* (*d*(**r, r**_*σ*_)), the total force at a position **r** exerted by all active morphodynamic processes at play within the tissue is

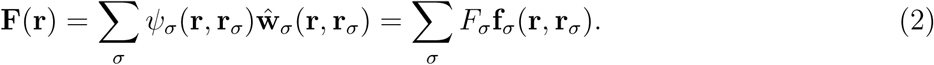

In what follows, to simplify notation, we often kept the dependence of **F**_*σ*_ on **r**_*σ*_ implicit and simply wrote **F**_*σ*_=**F**_*σ*_(**r**). The effect of the total force in Eq. (2) is also “felt” by dividing cells as it alters the mechanical state of their environment. Along its trajectory from **y**^(*i*)^ to **z**^(*i*)^ during cell division, a vertex at instantaneous position **p**^(*i*)^(*t*) will feel the force **F**(**p**^(*i*)^(*t*)) acting on it, on top of the viscous friction *η* of the local environment. This force exerts mechanical work on the vertex and alters the energetic cost of the dividing cell: the more intense the force **F**(**p**^(*i*)^(*t*)), the easier for the vertex to move along its direction. As a result, the long-range forces of Eq. (2) supply a part of the energy needed for shape changes during and after mitosis. The total change in a cell’s internal energy during division is thus due to the energy spent against friction forces plus the work performed by the external forces in Eq. (2). This can be quantified as

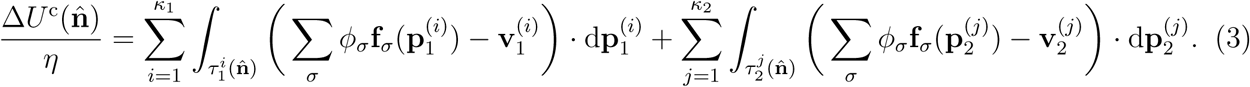

This is the energy that a dividing cell actively employs to change its shape. Δ*U*^c^ stems from the multiscale competition between the passive resistance of the local cellular neighbourhood (against which the dividing cell actively spends the energy given in Eq. (1)) and the long-range mechanical stress produced by the other concurrent active processes in Eq. (2), external to the dividing cell. Δ*U*^c^ is also a function of the cell’s division orientation through the trajectories 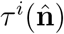. This competition between the viscous friction *η* of the local environment and the force **F**_*σ*_ generated by active morphological changes is captured by the ratio 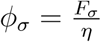. Be *v*_*m*_ a typical value of vertex displacement speed during cell division. Then, when *ϕ*_*σ*_ ≫ *v*_*m*_, the force exerted by the process *σ* dominates over the local resistance of the environment to deformation; on the other hand, when *ϕ*_*σ*_ ≪ *v*_*m*_, the force **F**_*σ*_ is negligible and viscosity dominates. This competition, captured by Eq. (3), is the cornerstone of our work and provides an energetic standpoint to understand energy fluxes between a dividing cell and its local and distant environments. Equation (3) quantifies the change in the internal energy of a dividing cell, as a function of the orientation of its mitotic plane. In the next Section, we will employ this equation to characterize the energetic behaviour of proliferating tissues in developing biological systems.

It is worth stressing at this point that the effect of mechanical forces in Eq. (3) is fully independent of the possible interphase strain they induce on the cell: in Eq. (3) the consequences of stress anisotropy are independent on whether the cell elongates during interphase in the direction of stress. This is indeed what was observed in several *in vivo* situations [34, 35]: mechanical stress in a tissue correlates with division orientations even in the absence of cellular interphase elongation in the direction of stress, leading to systematic violations of geometric rules for cell division.

## III. EMERGING BEHAVIOURS

### A. Isolated tissues

Equation (3) has several interesting consequences. For isolated tissues in the absence of mechanical cues (*ϕ*_*σ*_ = 0), cell geometry is expected to guide the orientation of cell division. The local part of Eq. (3), i.e., Eq. (1), is able to account for and physically explain this fact, as shown in Fig. 2 for 10^4^ cells, whose shape and vertices distribution were randomly simulated as explained in Section I of the Supporting Information. For each of them, Eq. (1) was numerically minimized with the help of biologically-relevant simplifying assumptions, as explained in details in Section II of the Supporting Information. As shown in Fig. 2, the energetically optimal direction 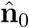 minimizing Eq. (1) correlates with both the main elongation axis 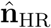 and the principal direction of the distribution of tricellular junctions (in our model, cell vertices) 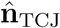. Interestingly, the agreement of Eq. (1) to 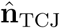 is higher than to 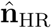, supporting tricellular junctions as the main geometrical cue for division orientation in 2D structures, as shown in [21]. This shows how the direction determined by cell elongation is often also the one along which shape changes can happen at the lowest energetic cost in viscous environments. Cells following HR (or the tricellular-junction rule) are the ones minimizing their energy loss.

This illustrates that cells following geometric rules have an energetic advantage.

### B. 2D tissue development: *Drosophila* dorsal thorax pupal epithelium

To further test the predictions of Eq. (1), we studied division orientation distribution during the development of the pupal epithelium of dorsal thorax of *D. melanogaster*. Using the dataset from [7], we reconstructed and tracked cells from the central patch of the developing tissue, during 14h of development. From this reconstruction, using the DRACO-STEM library [53], we further built a polygonal approximation of each cell which respects the original topology, as shown in Fig. 3, from which cell-junction (vertex) positions were extracted. As shown in [7], cell patches in the central zone of the pupal dorsal thorax develop with an isotropic, square-like geometry. This suggests that mechanical stress, if present, is highly isotropic in that zone. In turn, this means that the total external force in Eq. (2) acting at all positions in the reconstructed zone of the tissue is negligible and Eq. (1) should alone be able to recapitulate the observed division orientations.

The observed dynamics of shape change during and after cell division is highly variable. Both the daughter cell shape and the surface change after division do not show conserved patterns. We have nevertheless assumed that the “desired” shape of daughter cells is regular, and that deviations from this assumptions are due to strong mechanical interactions with neighbouring cells. To account for cell-to-cell variability, we optimized for each cell division the surface area ratio 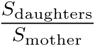 in a small range between 0.8 and 1.1: this range is what we most frequently observe in the dataset for the dynamics of the ratio 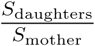 over few timepoints after division.

Excluding cells at the border of the reconstructed patch, to avoid reconstruction artefacts, left us with 3664 cell divisions. For each of them, we compared the division orientation with the main direction of the interphase distribution of cell-junctions, obtained by studying the vertex positions at two-third of each cell’s cell cycle duration. This geometry was also used to run the model of Eq. (1) as detailed in Section II of the Supporting Information. Specifically, a constant vertex speed *v* has been assumed. For comparison, we also run the model with non-negligible external forces in Eq. (3), assuming a a main direction of stress anisotropy along the y-axis of Fig. 3. Different choices of stress anisotropy direction led to similar results. Agreement results for both models are shown in Fig. 4.

The blue histograms in Fig. 4(a,b) show the distribution of angles between the actual division orientation and the cell-junction geometric rule. One finds a marked peak at low angles, with a low long-tail distribution at high angles, in accordance to what previously reported [21]. As expected, the best agreement of the model to observed orientations was obtained with Eq. (1), i.e., with no contribution of external forces or *ϕ*_*σ*_ = 0. Figure 4(a) shows the deviation angle distributions between the actual division orientation and the predictions of Eq. (1) (red), compared with the deviations from the main direction of cell-junction distribution (blue). Figure 4(b) shows the same comparison Eq. (3) and *ϕ*_*σ*_ = 10*v* (yellow). The model predicts no stress anisotropy to be present in the developing tissue. Both TCJ rule and the model with *ϕ*_*σ*_ = 0 work well in explaining the observed distribution of division directions, with average deviation angles of ∼29° for Eq. (1) and ∼30° for TCJ distribution. As expected from the results of subection III A, our model agrees with geometric rules (and in particular with cell-junction distribution) in absence of stress anisotropy. In addition, Eq. (1) predicts a higher fraction of orientations within 3° from the actual direction, and has fewer high-angle deviations (> 80°) than the distribution of TCJ. The result shows well that the model is able to infer the mechanical stress state of the tissue and to reproduce, and at the same time provide physical bases to, the correlation between tricellular junction positions and regulation of division orientation in epithelia, showing that cells follow the most energetically-favourable path during shape changes in mitosis.

### C. Multicellular structures under mechanical stress: a paradigm to understand embryonic morphogenesis

#### a. Ascidian epidermal morphogenesis during gastrulation

In many organisms, major morphogenetic processes are at play during key stages of development. These processes arguably produce markedly anistropic stress profiles in neighbouring tissues. Such is the case in developing animal embryos, during the morphogenetic process known as gastrulation [54]. Figure 5(a) illustrates the process of gastrulation in embryos of the ascidian *P. mammillata*, a model organism to study animal morphogenesis [55]. Around 4 hours after fertilization, these embryos undergo major shape change processes, collectively known as gastrulation, which begin with the invagination of a group of cells [56], whose active shape changes drive the whole process as schematically depicted in Fig. 5(a) for control (Ctrl) conditions (see also Fig. 2 of the Supporting Information). Part of the epidermis of these embryos is at one-to few-cell distance from the gastrulating core and is hence strongly subjected to the induced mechanical stress. Interestingly, during these processes no cell migration nor cell death is known to happen.

**FIG. 5:**
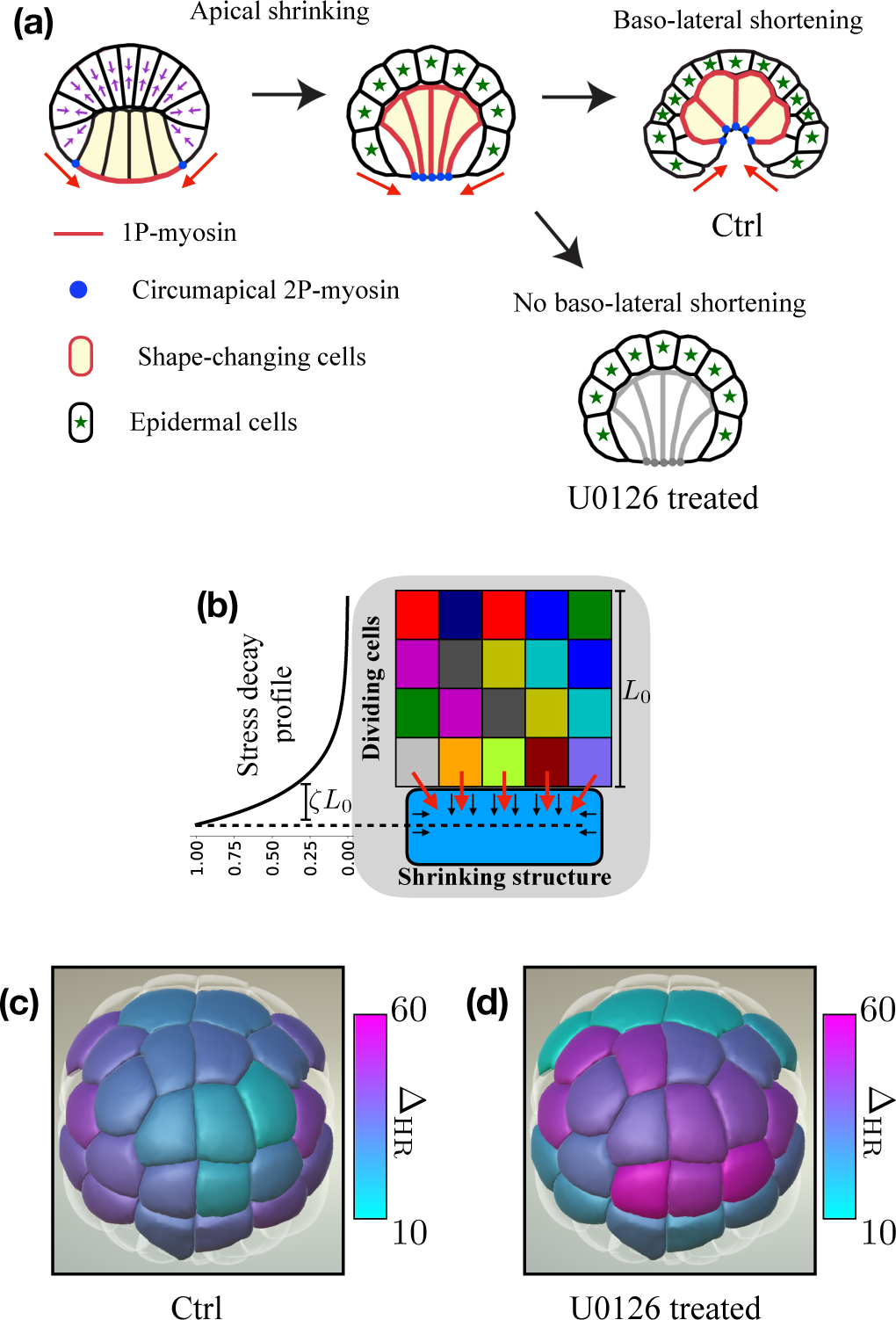
**(a)**: Gastrulation in embryos of *P. mammillata* in both Control and U0126 treatment conditions (adapted from [56]). Panel **(b)**: system of 20 cells, of characteristic linear dimension *L*_0_, in contact with a shrinking structure (panel **(a)**), with exponentially-decaying stress profile of length scale *ζL*_0_. This serves as a model to study the morphogenesis of the epidermis of embryos of *P. mammillata* during gastrulation. Thanks to the invariant lineage of ascidian embryos, homologous cells can be identified in different embryos. Panels **(c)** (for Control) and **(d)** (for U0126 treatment) show the average angular deviation Δ_HR_ (in degree) from HR for three rounds of divisions in ascidian epidermis. The average for each cell is taken over all divisions and across two embryonic datasets for Control conditions (taken from [19]) or one dataset for U0126 treatment and visualised through the morphological browser MorphoNet [45]. To facilitate comparison and exploiting the invariant lineage of ascidian embryos, the deviations in panels **(c)**-**(d)** are projected onto the same wild-type embryo reconstruction.

To evaluate the role of mechanical stress in epidermal morphogenesis, we also partly blocked gastrulation by FGF/ERK signalling inhibition (U0126 inhibitor). Affecting the balance between FGF and Eph signaling was previously shown to impair gastrulation [57] by a fate switch of A-line endoderm precursors to trunk lateral cells [58]. Embryo treated with U0126 would undergo the first of the two sequential processes of gastrulation (apical shrinking) but would not proceed through the second one (baso-lateral shortening), as depicted in Fig. 5(a) for U0126 treatment and as shown in Fig. 2 of the Supporting Information. This arguably produces a weaker stress profile propagating to epidermal precursors, since the contribution to mechanical forces of baso-lateral shortening is missing. The morphogenesis of embryonic ascidian epidermis is a quasi-2D process, in spite of the global 3D curved structure of the developing embryo (Section IV and Fig. 3 of Supporting Information).

To study the consequences of the competition between local viscosity and long-range propagation or transmission of mechanical forces in this system, we analysed the theoretical situation depicted in Fig. 5(b) in which one shrinking structure (in blue) (representing for instance a group of cells) exerts a force *F*_0_**f**_0_ on a simulated system initially composed of *n* cells, with characteristic linear dimension *L*_0_, undergoing three rounds of cell divisions.

Thanks to the transparency of *P. mammillata* embryos and to state-of-the-art 3D reconstruction of live embryonic development, high-quality 3D dynamic data on their development is now available [19]. Remarkably, ascidian embryos develop with a fixed lineage, which allows to identify homologous cells across several different embryos. We have hence calculated, as detailed in Section III of the Supporting Information, the angular deviation Δ_HR_ of division orientation from HR for each epidermal cell across three rounds of cell divisions in two different control embryos and in the U0126-treated embryo. The average value across all divisions within each clone (i.e., the set of progeny of a single initial cell) and across the two control embryos is shown in Fig. 5(c), projected onto the 22 cells at the 64-cell stage which will give rise to epidermis. A pattern of deviations emerges, for cells mostly localised around the tissue border.

We performed the same analysis on the U0126-treated embryo. The average value per clone is reported in Fig. 5(d), represented on the same embryo used in Fig. 5(c) to facilitate comparison. The deviations observed in Ctrl conditions at the interface between epidermis and mesendodermal tissues are far less present in the U0126 treated embryo: affecting the stress profile induced by gastrulation seems to have an impact on the division orientation of epidermal cells.

These observations make the epidermal morphogenesis of *P. mammillata* an ideal system to test our theory.

#### b. Application of the model to ascidian epidermis

We hence set out to study the toy model of Fig. 5(b) with different magnitudes of external mechanical forces, in order to understand whether the division orientation and the global shapes thus produced can explain the phenomenology of ascidian epidermis. Orientation of each cell division is determined according to our model, hence minimizing Eq. (3) as explained in Section II of the Supporting Information. After division, due to the concurrent changes in cell shape, the total tissue is out of mechanical equilibrium. This produces, during interphase, a dynamics of relaxation towards the tissue rest shape, which comes from the competition between interacting cells trying to minimize their configurational energy. Under normal conditions, this energy for an isolated cell is minimized for spherical shape due to the cell’s cortical tension [59]. The dynamics of a 2D tissue has largely been studied and is well captured by the many-cell energy function [46]

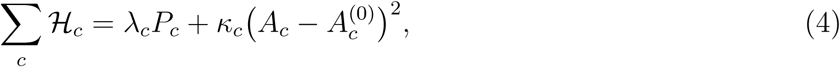

in terms of cortical tension *λ*_*c*_ along the cells perimeter *P*_*c*_ and surface elasticity *κ*_*c*_ of the cells area *A*_*c*_ around an equilibrium surface 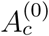. The latter accounts at once for volume compressibility and for the actomyosin-mediated elastic properties of a cell surface. The final rest shape is achieved as the minimum of the multicellular energy in Eq. (4).

Viscous forces during cell division depend on the speed at which each vertex trajectory is travelled. To keep things as simple as possible, we assume here a constant vertex displacement speed *ν*. Recalling the definition of the parameters 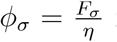 in Eq. (3), one can introduce the dimensionless version of *ϕ*_*σ*_ as the order parameter 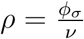, measuring the relative intensity of external to viscous forces: *ρ* ≫ 1 means dominating contribution of external forces. Note that *ρ* can also be seen as a measure of the amount of anisotropy of mechanical stress compared to the viscosity of the medium. This is because an isotropic mechanical stress would imply that each direction of vertex movement is equally affected by long-range forces and hence no bias in the choice of optimal division direction would be produced. At the same time, one expects that the effect of localized morphogenetic processes is not perceived with the same intensity everywhere in a tissue. This means that the stress profile induced by each of these processes decays with the distance from the position of the process itself, as described by the function *h* (*d*(**r, r**_*σ*_)) in Eq. (2). Stress decay is assumed here to be an exponential function 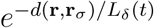, with a time-dependent length-scale *L*_*δ*_(*t*) = *δ*(*t*)*L*_*d*_ (*δ*(*t*) = 1 for the first two divisions and *δ*(*t*) = 2 for the last). This time-dependence account both for the increasing density of cell junctions with cell divisions, known to help propagating tissue-scale stress profiles [60]. The fractional stress decay lengthscale is *ζ* = *L*_*d*_*/L*_0_. Between divisions, the system evolves towards its interphase equilibrium according to the Hamiltonian in Eq. (4), with parameters initially fixed at 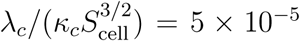 (*S*_cell_ being the surface of each of the 20 initial cells), to account for cell volume incompressibility. We will however show that the results of the model are highly robust against noise in this ratio.

We characterize the dynamics of cell divisions by two main quantities: how much each cell deviates from the predictions of pure cell geometrical rules, and how aligned consecutive divisions within the same clone are. The latter quantity gives information on whether cells tend to orient their division plane along a fixed axis, independently on their shape. This quantity is also correlated to deviations from geometrical rules: it is known that the agreement to geometrical rules often gives rise to alternating orthogonal directions for division orientations within one clone [27]. We measure here the cellular elongation as the main direction of the distribution of its vertices, since cell shape itself is defined by its vertices in the vertex-based model we employ. At the same time, this is biologically relevant as it corresponds to the main direction of tricellular junction distribution, as already commented in previous Sections. The first quantity is measured by the parameter *α =* 1 − ⟨𝒳_CJ_⟩/90, with 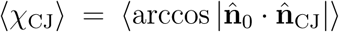 being the average angle between the direction of division and the main direction of distribution of tricellular junctions. The average is taken over every cell at each round of division. *α* is 1 for perfectly aligned and 0 for fully orthogonal predictions.

To define a measure for the second quantity (the alignment of consecutive divisions within a clone) consider, for a set of initial cells 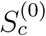, the parameter

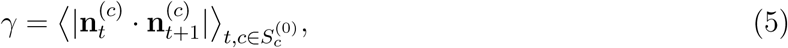

measuring the correlation between the orientations **n** of any two consecutive divisions at *t* and *t* + 1 within the same clone of any cell 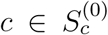. The average is taken over all cells and all pairs of consecutive division rounds.

We use these two parameters to study the influence of external forces on the initial set of 20 cells of Fig. 5(b). Based on what we observe in ascidian epidermis (see Sections III and IV of Supporting Information and results in [19]), we assume that a dividing cell increases its surface area by a factor 2^1*/*3^ (representative of what measured during embryonic development, as shown in Section IV of the Supporting Information) and that daughter cells, after divisions, tend to acquire regular shapes. The left plot of Fig. 6(a) shows *α* as a function of *ρ* (ratio of intensity of active to viscous forces) and *ζ* (penetration length of mechanical stress in the tissue). Two phases appear in the response of the system: as expected at low *ρ* (dominant viscosity), Eq. (3) correlates very well with individual-cell geometry and *α* reaches high values; increasing *ρ*, however, the system undergoes a behavioral phase transition of the continuous type at a critical value *ρ*_*c*_ of the order parameter. As shown in Fig. 6(a), this critical value is only weakly dependent on *ζ*, provided *ζ* itself is above a critical value *ζ*_*c*_. At high *ρ* a new phase emerges in which the agreement between Eq. (3) and geometrical rules is rapidly lost (*α* sharply decreases). To understand the cause of this phase transition we looked at the correlations between cells within the simulated tissue.

**FIG. 6:**
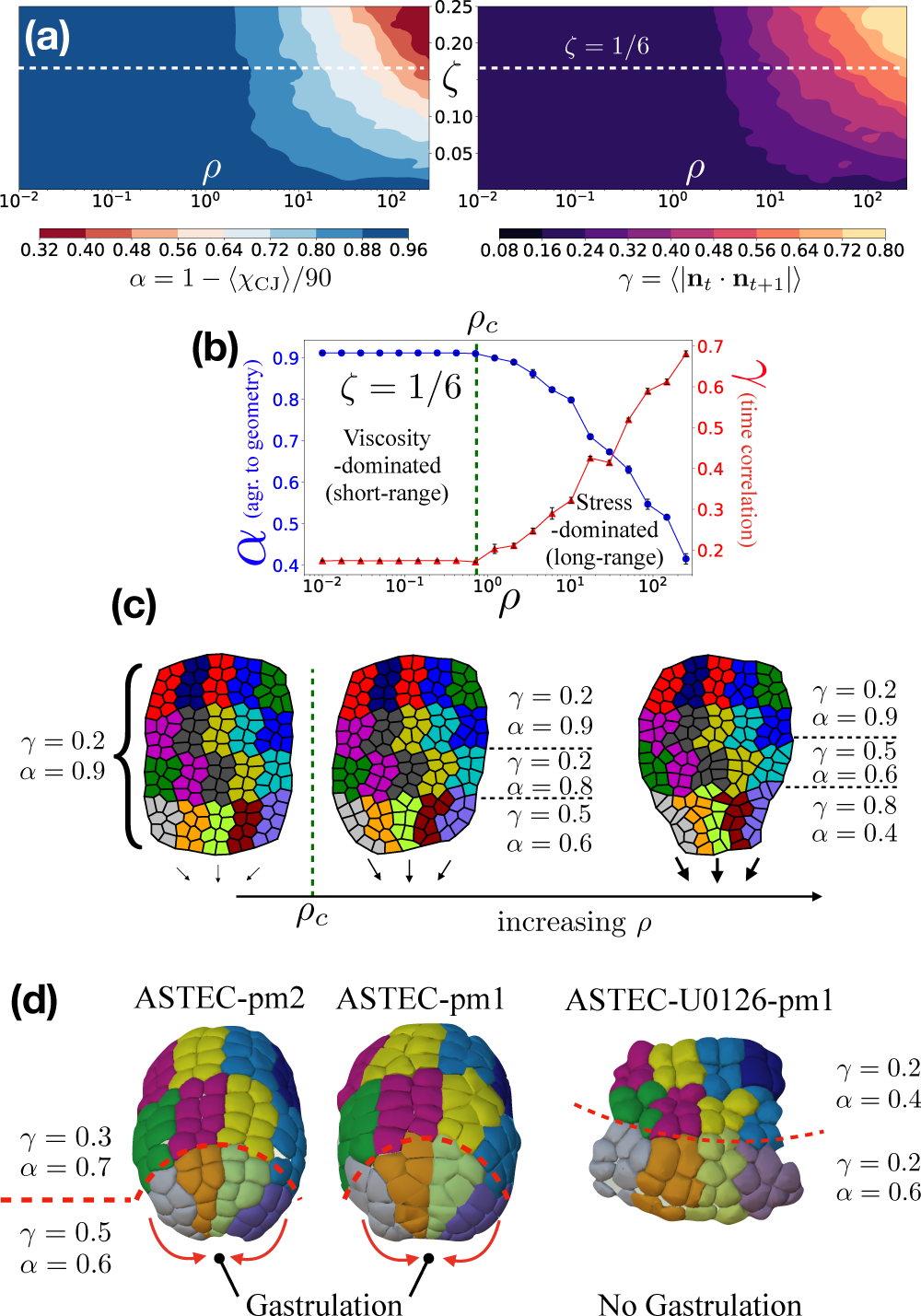
Panels **(a)** and **(b)** show the predictions of Eq. (3) for the simulated system of Fig. 5(b). A marked transition is found between two phases, (panel **(a)**), for both the agreement of Eq. (3) to geometrical rules *α* and the average time-correlation of division orientations in clones *γ*, as a function of the order parameters *ρ* and *ζ*. The first phase, (viscosity-dominated, low *ρ*) shows a high agreement *α* to the geometrical rules for cell division and a low time-correlation parameter *γ*. The second phase (stress-dominated, high *ρ*) shows poor agreement to geometrical rules and high temporal correlations. The transition between the two is continuous, as seen in panel **(b)**, showing the transition at fixed stress fractional penetration length *ζ* = 1*/*6 (*γ* in red triangles, right vertical scale; *α* in blue dots, left vertical scale). Black vertical bars give the standard deviations for 20 randomly simulated values of the ratio 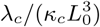. Panel **(c)** shows the consequences of this transition on the global tissue shape: above the critical value *ρ*_*c*_, the shape of the whole tissue start being affected by the externally-induced mechanical stimuli. This is a consequence of the time correlations of division orientations, building up in the tissue due to long-range mechanical cues. The dynamics of stress propagation can be clearly read in the shape of the clones (collection of all progeny) of each of the 20 initial cells. These predictions were tested in the development of the epidermis of embryos of *P. mammillata* (panel **(d)**), in both control condition (ASTEC-Pm1, ASTEC-Pm2) and U0126 treatment. Due to the force produced by shape changes of invaginating cells during gastrulation, Ctrl embryos show the same transition in clones shape and cellular distribution as the one predicted by the model at intensities *ρ* slight past the critical value. The U0126-treated epidermis shows no such transition, in agreement with the prediction of the model at low *ρ* values due to the absence of baso-lateral shortening. Control-conditions embryonic 3D reconstruction from [19].

Figure 6(a) (right) and 6(b) show that the transition to the new phase in the tissue is correlated to the appearance of time-correlations within each clone, as measured by the parameter *γ* [Eq. (5)]. These time-correlations are in turn induced by the stress dynamics within the tissue.

To test the influence of mechanical relaxation parameters of Eq. (4), simulations at fixed *ζ* = 1*/*6 were run with random values for 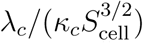, drawn out of a Gaussian distribution centered around 5 × 10^−5^ and of standard deviation 10^−5^. Figure 6(b) shows, for each simulated value of *ρ*, the average and the standard deviation (in black vertical bars) for 20 random values of 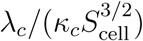.

The onset of time correlations *γ* past the critical value *ρ*_*c*_ suggests that the shape of clones may be strongly altered in the presence of external forces, even without single-cell elongation in the direction of these forces, simply due to time correlations between division orientations and shape changes during cytokinesis. This is indeed the case, as shown in Fig. 6(c): past the critical value *ρ*_*c*_, clones start showing a more elongated form, tending to a linear shape at high *ρ* values. This in turn affects the global tissue shape, which tends to follow the change in shape of the external shrinking structure. The two phases can coexist in the same tissue, as shown in the rightmost example of 6(c), in which the cells facing away from the external force are in the viscosity-dominated phase and follow geometrical rules (*α* = 0.9, *γ* = 0.2), while the ones directly facing the force are in a stress-dominated phase with high temporal correlations and high deviations from geometry (*α* = 0.4, *γ* = 0.8).

Thanks to the availability of several 3D reconstructions of live embryonic development [19], we analysed division orientations and apical geometry of epidermal cells in two embryos of *P. mammillata* (see also Fig. 2 of the Supporting Information): the coexistence of these two phases is observed *in vivo*, and elongated epidermal clones are built up while following the active shape changes of gastrulation as exemplified in Fig. 6(d) for the two Control embryos ASTEC-Pm1 and ASTEC-Pm2. The cellular distribution in each clone observed in this process is the same as the one predicted by Eq. (3). In addition, the overall tissue shape shows the same structure as the high-stress tissues of Fig. 6(e), with a similar phase transition marked by a change in *α* from 0.7 ± 0.01 to 0.6 ± 0.03 and in *γ* from 0.3 ± 0.03 to 0.51 ± 0.04.

On the other hand, such transition is much weaker in the U0126-treated embryo ASTEC-U0126-Pm1, in which gastrulation is largely blocked (rightmost embryo in Fig. 6(d)): the value of *γ* is constant everywhere in the analysed epidermis and the clones shape is everywhere similar to what expected at low *ρ* (weak external forces). The overall shape of the tissue, in spite of some noise due to the marked change in global embryonic geometry (Fig. 2 of Supporting Information), closely resembles what predicted by our model before the behavioural transition (leftmost simulation in Fig. 6(c)), where clones have rectangular shapes and *γ* = 0.2 everywhere: this is exactly what we observed in the U0126-treated embryo.

## IV. CONCLUSIONS

In this work, we propose a mechanism by which developing biological structures maintain a global shape consistency in spite of major local changes in their form. Through a reformulation of the process of cytokinesis in terms of its energetic cost, we show that cells can read shape dynamics of their local and remote environments, and react by orienting their division in a way that minimizes the work spent against the passive resistance of their surrounding. The cell main geometrical axis emerges as the direction along which the work performed by a cell during division and against the passive resistance of its local environment is minimal. Which mechanisms are at play for a cell to be able to choose the “path of least resistance” for its division is an open question.

Division orientation is determined by the position and orientation of mitotic spindle. These, in turn, are known to be determined by forces exerted by microtubules as they interact with the cell cortex. In our view, the mechanisms guiding a cell to its energetically-favourable trajectories during mitosis are able to bias these forces following external mechanical cues.

As shown in a number of recent works [61, 62], cells are able to sense the mechanical state of their environment by mechanosensing pathways. Specifically during cell division, one of these pathways involves the state of tension of astral microtubules [36], which is affected by how easily the dividing cell can deform its local environment. The cell could thus probe a number of division orientations to identify in which direction small deformations produce the optimal tension on the spindle-assembly microtubules.

We remark that cells dividing along energetically-optimal directions have an advantage over the ones who do not. The complex machinery of spindle positioning might have been developed under such evolutionary pressure. In *D. melanogaster* [21], the accumulation of certain proteins at three-cellular junctions is used by the cell to orient its spindle poles. As we have shown in this work, in the absence of external stress such orientation correlates with the energetically-optimal direction minimising Eq. (1). We can further suppose that a state of mechanical stress can bias the distribution of such proteins and, hence, influence the division orientation reproducing the predictions of Eq. (3). These proteins might be the mechanisms selected by evolution because energetically favourable.

When applying our theory to the development of the pupal epithelium of dorsal thorax of *D. melanogaster*, we were able to infer the absence of stress anisotropy in the tissue and to recapitulate the known rule for division orientation based on interphase distribution of tricellular junctions [21].

However active processes of shape dynamics can bias such energetic cost and provide multiscale cues to guide cell division, in agreement with what is observed, for instance, in the *Arabidopsis* meristem [33]. In these cases, reproducible and systematic deviations from long-axis rules emerge, as repeatedly experimentally reported [33–35]. We provide a theoretical framework to explain and understand these deviations: the competition between local and long-range stimuli gives rise to a transition between two distinct phases of a multicellular system, marked by the onset of time-correlations within cellular clones. Remarkably the co-existence of these two phases, predicted by our model, is found *in vivo* in the epidermis of the developing embryo of the ascidian *P. mammillata*. Our model is further able to account for the phenotype of ascidian embryos after FGF/ERK signalling inhibition, which partially inhibits endoderm invagination and as a consequence changes the orientation of cell divisions in epidermis.

Our model can thus explain both the known geometrical rules for cell division and the systematic deviations from them observed in certain developmental contexts. We note however that, due to the lack of shape changes during plant cell division, the extension of this framework to vegetal cells is yet an open question and deserves further analysis. Both geometrical rules and deviations emerge as particular cases of a more general rule, which is the consequences of an energy exchange between a dividing cell and its environment. This exchange provides a way by which cells can sense and react to external mechanical forces, fully independent of the presence of interphase cell elongation in the direction of these forces. In this way, tissue deformation and shape dynamics can be produced without the need of individual cell deformations. Our result sheds new light on the physics of morphogenesis and on the mutual interplay between morphodynamics and mitosis, and suggest that oriented cell division is one of the contributing mechanisms allowing long-range transfer of information in biological structures. Such information transfer, under the form of internal mechanical stress, can be read in the shape of cellular clones within a proliferating tissue, providing a new paradigm to infer and follow force dynamics in living systems.

## Supporting information

*Section I.* **Random simulations of dividing cells.** The section provides details on the model applied to randomly simulated cells in Fig. 2.

*Section II.* **Implementation of the optimal division model.** The section provides details on the mathematical and numerical implementation of the models given by Eqs. (1) and (3).

*Figure 1.* **Scheme of cell division process.** Schematic representation of the simulated process of cell division.

*Section III.* ***P. mammillata* embryonic epidermal cells and analysis of their division directions.** The section provides details on how epidermal cell divisions in the datasets of ascidian embryonic development have been analysed.

*Figure 2.* **Reconstruction of gastrulating embryos of *P. mammillata*.** Reconstructed ascidian embryos before, during and after gastrulation, both in WT conditions and after U0126 treatment.

*Section IV.* **Geometrical properties of epidermal precursors in developing ascidian embryos.** The section provides details on the geometry of epidermal cells in ascidian embryos. Specifically, their clonal shapes, their apical surface and their volumes are analysed.

*Figure 3.* **Departure from planar geometry for each clone of epidermal precursors.**

*Figure 4.* **Schematic view of apical surface changes after cell division in ascidians.**

*Figure 5.* **Dynamics of the average elongation of apical surface of epidermal cells in ascidian embryos.**

*Figure 6.* **Dynamics of the average relative variation of the elongation of apical surface of epidermal cells in ascidian embryos.**

*Figure 7.* **Distribution of sister cells volume ratio in ascidian epidermis.**

*Section V.* **U0126-treated *P. mammillata* embryo and its phenotype.** The section provides details on the U0126 treatment of ascidian embryos.

*Table 1.* **Parameters and variables of the model.**

## ACKNOWLEDGMENTS

PL and EF were members of CNRS, CG was a member of INRIA. Work in PL and CG’s teams was funded by the Dig-Em project (ANR-14-CE11-0013-01) and the Institut de Biologie Computationnelle of Montpellier (IBC; ANR-11-BINF-0002). JL was supported by Dig-Em, BL by Dig-Em and IMPULSION project Mecafield from IDEXLYON. The authors wish to thank Dr. G. Cerutti for his help in reconstructing the *Drosophila* dataset; Dr. L. Guignard and Dr. U.-M. Fiuza for their outstanding efforts in making 3D reconstructions of ascidian embryos available; Dr. Y. Bellaïche, Dr. J. Traas, Dr. O. Hamant, Dr. O. Ali and Dr. L. Sundermann for carefully reading the manuscript and for fruitful discussions. They also wish to thank Dr. B. Guirao and Dr. Y. Bellaïche for generously sharing the dataset of *Drosophila* dorsal thorax.

All authors contributed to the conception of the original idea. BL developed the theory and the physical model, performed simulations, analysed datasets and results and wrote the paper; JL and EF contributed to the development of the theory and its connection to ascidian embryos; JL prepared datasets for analysis and performed the U0126 treatment of ascidian embryos; EF contributed to the analysis of the *Drosophila* dataset; PL and CG contributed to the development of the theory, the interpretation of results and the writing of the paper. All authors contributed to the discussion and interpretation of the main results.

