## Supporting Information for "Multiscale mechanical model for cell division orientation in developing biological systems"

### I. RANDOM SIMULATIONS OF DIVIDING CELLS

To test the agreement between Eq. (1) of the main manuscript and either Hertwig's rule or the tri-cellular junctions distribution, we have studied divisions for $10^4$ randomly simulated cells. The geometry and vertex distribution was randomly chosen as follows:

- the shape of each cell is determined by the perimeter of an ellipse, whose eccentricity and main axis orientation were randomly chosen. For the eccentricity, values were randomly drawn out of a uniform distribution over the interval $[0.2, 1)$ in order to avoid shapes without clear geometrical cues. Each cell was then determined as a collection of vertices lying along the perimeter of this ellipse;
- the in-plane orientation of each ellipse's major axis, which gives also the cell's long axis, was randomly simulated by drawing two numbers $x_v$ and $y_v$ out of a gaussian distribution of zero mean and unit variance and using then the vector $\mathbf{n}_a = \frac{1}{|(x_v,y_v)|}(x_v, y_v)$;
- for each cell, the number of vertices was randomly chosen from the interval $[4, 50]$ with uniform probability. Although cells with more than 10 vertices are extremely rare in living tissues, we have pushed the limits in our simulations to 50 to show the robustness of our model against vertex number;
- among the vertices constituting each cell, a randomly drawn subset was positioned at irregular angles, while the rest at the expected positions of a regular angular distribution;
- the angular position of each irregular vertex was drawn at random from a Gaussian distribution, centred on the corresponding regular angular position and with constant standard deviation $\sigma_v = 0.3\,\text{rad}$;

For each cell this, the main direction of tri-cellular junctions was calculated by a principal-component analysis of its vertices distribution, while the main elongation was identified as the direction of the ellipse's main axis.

Finally, Eq. (1) of the main text was minimised to find the energetically-optimal division direction, as detailed in the next section.

### II. IMPLEMENTATION OF THE OPTIMAL DIVISION MODEL

Here we detail the technical aspects of the implementation of the optimal division model in Eq. (1) of the main text. The main difficulty is to minimize the dissipation

$$W_\eta = \eta \sum_{i=1}^{\kappa_1} \int_{\tau_1^i(\hat{\mathbf{n}})} \mathbf{v}_1^{(i)} \cdot \mathrm{d}\mathbf{p}_1^{(i)} + \sum_{j=1}^{\kappa_2} \int_{\tau_2^j(\hat{\mathbf{n}})} \mathbf{v}_2^{(j)} \cdot \mathrm{d}\mathbf{p}_2^{(j)}, \tag{1}$$

as a function of the set of trajectories $\{\tau_1^i\}$ and $\{\tau_1^j\}$ followed, respectively, by the vertices which will belong to daughter 1 or daughter 2 during shape dynamics of anaphase and cytokinesis. These trajectories, given a fixed initial

---

† Current address: Institut Clément Ader, Université Toulouse III, CNRS, INSA, ISAE-SUPAERO, Mines-Albi, France

shape of the dividing cell, depend on three main quantities: the velocity of each vertex during shape dynamics, the position and orientation of division plane, and the final shape of the daughter cells.

To simplify the theoretical and numerical treatments, we employ the following biologically-motivated assumptions:

1. The trajectory travelled by each vertex in the dividing cell is linear;
2. The speed at which these trajectories are travelled is constant in time and the same for all vertices (hence we neglect the transient regimes of onset and end of the vertex movement);
3. The division plane passes through the geometric center of the apical surface of the dividing cell, and daughter cells immediately after division have the same apical area $K_a S_m$, where $S_m$ is the apical area of the mother cell and $K_a > 0$;
4. The final shape of the apical surface of daughter cells tends to be round. The final shape of the vertices of each daughter cell (i.e., the shape they would acquire if the resistance of the surrounding environment was negligible) is thus a regular polygon.

The experimental evidence justifying assumptions 3 and 4 above is provided in Sections IVB and IVC of this Supporting Information.

Under these hypotheses, each term $\int_{\mathcal{T}_1^i(\hat{\mathbf{n}})} \mathbf{v}_1^{(i)} \cdot \mathrm{d}\mathbf{p}_1^{(i)}$ in Eq. (1) can be given a much simpler expression.

Be $\mathbf{p}_i$ and $\mathbf{p}_f$ the initial and final positions of a specific vertex along its trajectory. Due to assumptions 1 and 2, one can write its velocity along the trajectory as

$$\mathbf{v} = v \frac{\mathbf{p}_f - \mathbf{p}_i}{|\mathbf{p}_f - \mathbf{p}_i|} \tag{2}$$

and the position of the vertex in time becomes

$$\mathbf{p}(t) = \mathbf{p}_i + vt \frac{\mathbf{p}_f - \mathbf{p}_i}{|\mathbf{p}_f - \mathbf{p}_i|}. \tag{3}$$

The trajectory of the vertex is thus Eq. (3) travelled from time $t_i = 0$ to time $t_f = \frac{|\mathbf{p}_f - \mathbf{p}_i|}{v}$.

The work of viscous forces of the vertex along this trajectory thus reads

$$w_\eta = \eta v \int_{\mathcal{T}} \frac{\mathbf{p}_f - \mathbf{p}_i}{|\mathbf{p}_f - \mathbf{p}_i|} \cdot \mathrm{d}\mathbf{p} = \eta v^2 \int_0^{\frac{|\mathbf{p}_f - \mathbf{p}_i|}{v}} \frac{\mathbf{p}_f - \mathbf{p}_i}{|\mathbf{p}_f - \mathbf{p}_i|} \cdot \frac{\mathbf{p}_f - \mathbf{p}_i}{|\mathbf{p}_f - \mathbf{p}_i|} \mathrm{d}t = \eta v |\mathbf{p}_f - \mathbf{p}_i|, \tag{4}$$

i.e., it becomes a function of only the distance between the initial and final position of the vertex.

Of course, while the initial position $\mathbf{p}_i$ of the vertex is fixed by the interphase geometry of the mother cell, its final position $\mathbf{p}_f = \mathbf{p}_f(\hat{\mathbf{n}}, \ldots)$ is a function of the division direction (hence, the direction of the plane separating the two daughter cells) and of additional parameters specifying the final geometry of the daughter cell the vertex will belong to.

Given a mother cell with $N_m$ vertices and $S_m$ apical area, thanks to the assumption 3 a specific direction $\hat{\mathbf{n}}$ of the mitotic plane unambiguously determines two subsets $s_1$ and $s_2$ of, respectively, $N_1(\hat{\mathbf{n}})$ and $N_m - N_1(\hat{\mathbf{n}})$ vertices, which represent the vertices of the mother inherited by, respectively, daughter 1 and daughter 2. Adding to both these sets the two new vertices $\mathbf{u}_0(\hat{\mathbf{n}}), \mathbf{u}_1(\hat{\mathbf{n}})$ created by the interface separating the two daughters, one has two sets $d_1$ and $d_2$ of, respectively, $\kappa_1(\hat{\mathbf{n}}) = N_1(\hat{\mathbf{n}}) + 2$ and $\kappa_2(\hat{\mathbf{n}}) = N_m - N_1(\hat{\mathbf{n}}) + 2$ elements. See Fig. 1(a) for a schematic representation in the case $N_m = 5$. The sets $d_1$ and $d_2$ contain the initial positions of all vertices.

Using assumption 4, the final vertex positions are also specified as the points of a regular polygon of area $K_a S_m$ and with a side parallel to $\mathbf{u}_0(\hat{\mathbf{n}}) - \mathbf{u}_1(\hat{\mathbf{n}})$. This uniquely determines two new sets $g_1$ and $g_2$ of $\kappa_1$ and $\kappa_2$ vertices, specifying the final positions of each vertex along its trajectory.

Since, however, the edge $\mathbf{u}_0 - \mathbf{u}_1$ must be shared between the two daughters, there can be situations in which it is not possible to respect at once assumptions 3 and 4. Such situations arise because the shared edge $\mathbf{u}_0 - \mathbf{u}_1$ fixes the length of the side of each regular polygon, and regular polygons with different number of vertices having the same edge length have different areas. When these cases arise, we relax assumption 3 and simulate two regular polygons with edge length $\frac{l_1 + l_2}{2}$, $l_1$ ($l_2$) being the edge length of a regular polygon with $\kappa_1$ ($\kappa_2$) vertices and surface area $K_a S_m$.

Summarizing, specifying $\hat{\mathbf{n}}$ entirely determines the set of initial and final points for each vertex, from which Eq. (4) and, hence, Eq. (1), can be directly calculated.

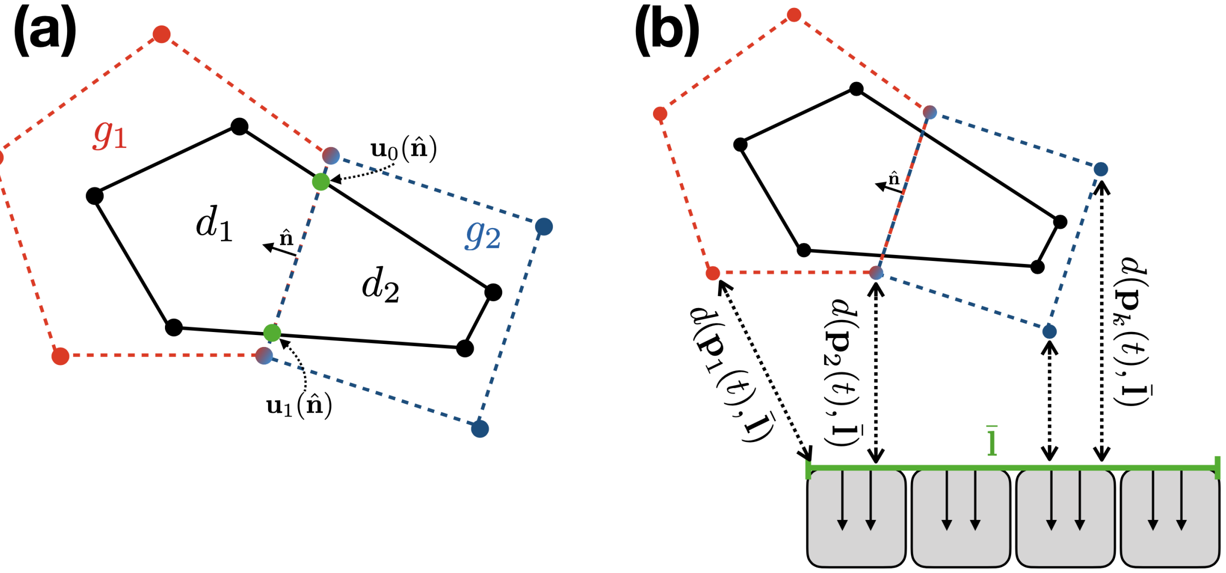


FIG. 1: **(a)**: Schematic representation of the simulated process of cell division. A mother cell with 5 vertices divides along the direction $\hat{\mathbf{n}}$, giving rise to two daughters with, respectively, 5 and 4 vertices. **(b)**: Representation of gastrulating cells (grey cells) used for the simulations of ascidian epidermal morphogenesis.

The minimisation of Eq. (1) with $\hat{\mathbf{n}}$ is performed numerically for each cell, by sampling 500 equally spaced directions in the plane and calculating Eq. (1) for each of them. The optimal direction is then chosen as the one producing the lowest value of $W_\eta$.

The strategy to minimise Eq. (3) of the main manuscript follows the same line, the only difference being the fact that the work of active forces $\mathbf{F}(\mathbf{r})$ must also be calculated. Since trajectories are assumed to be linear and active forces are kept constant during each division, the integral $\int_\tau \mathbf{F}(\mathbf{p})\mathrm{d}\mathbf{p}$ can easily be evaluated numerically by sampling the function $|\mathbf{F}(\mathbf{p}(t))|\cos(\theta(t))$ over 100 points in time, uniformly distributed in the interval $[0, \frac{|\mathbf{p}_f - \mathbf{p}_i|}{v}]$. Here $\mathbf{p}(t)$ is the point trajectory given in Eq. (3) and $\theta(t)$ is the angle between $\mathbf{p}_f - \mathbf{p}_i$ and $\mathbf{F}(\mathbf{p}(t))$.

As explained in the main text

$$\mathbf{F}(\mathbf{r}) = \sum_\sigma \mathbf{F}_\sigma(\mathbf{r}, \mathbf{r}_\sigma) = \sum_\sigma F_\sigma h\big(d(\mathbf{r}, \mathbf{r}_\sigma)\big)\hat{\mathbf{w}}_\sigma(\mathbf{r}, \mathbf{r}_\sigma). \tag{5}$$

For the simulations in Figure 6 of the main text, we have considered one external active process only, which means that $\mathbf{F}(\mathbf{r}) = Fh\big(d(\mathbf{r}, \mathbf{r}_0)\big)\hat{\mathbf{w}}(\mathbf{r}, \mathbf{r}_0)$, where $\mathbf{r}_0$ indicates the position of the external active process. In particular, we have considered a row of shrinking cells as represented in Fig. 1(b). The vector distance of each point $\mathbf{p}(t)$ along a vertex trajectory to the external active process is thus being defined as the vector between $\mathbf{p}(t)$ and the closest point $\mathbf{r}_0(\mathbf{p}(t))$ on the green segment $\bar{\mathbf{l}}$ of length $l$ in Fig. 1(b), representing the side of the shrinking cells facing the dividing cell. In the simulations for Figure 6 of the main text, the length $l$ is chosen to be the same as the length of the side of the initial 20-cell system facing the shrinking row.

Finally, the magnitude of the force in Eq. (5) becomes

$$|\mathbf{F}(\mathbf{p}(t))| = Fh\big(d(\mathbf{p}(t), \bar{\mathbf{l}})\big), \tag{6}$$

$d(\mathbf{p}(t), \bar{\mathbf{l}})$ being the point-segment distance.

We also stress here that, by construction, the energies in Eqs. (1) and (3) of the main text increase with increasing number of vertices. This, however, does not lead to any inconsistencies as long as one does not compare values of energy dissipation of two different cells. The procedure we have detailed here simply consists in comparing the values of Eqs. (1) and (3) calculated for the same cell (hence, at fixed number of vertices) along different possible division orientations $\hat{\mathbf{n}}$. We are not interested in the actual value of this dissipation, which depends on the specific model for cell geometry and mechanics, but rather in the position of its minimum. Hence, the scaling of Eqs. (1) and (3) with vertex number does not pose any problems in our model.

However, if one wants to compare the dissipation for two different cells, one should calculate the average dissipation per vertex simply by dividing the two terms in Eqs. (1) and (3) (one term per daughter cell) by the number of vertices of the respective daughter cell ($\kappa_1$ and $\kappa_2$).

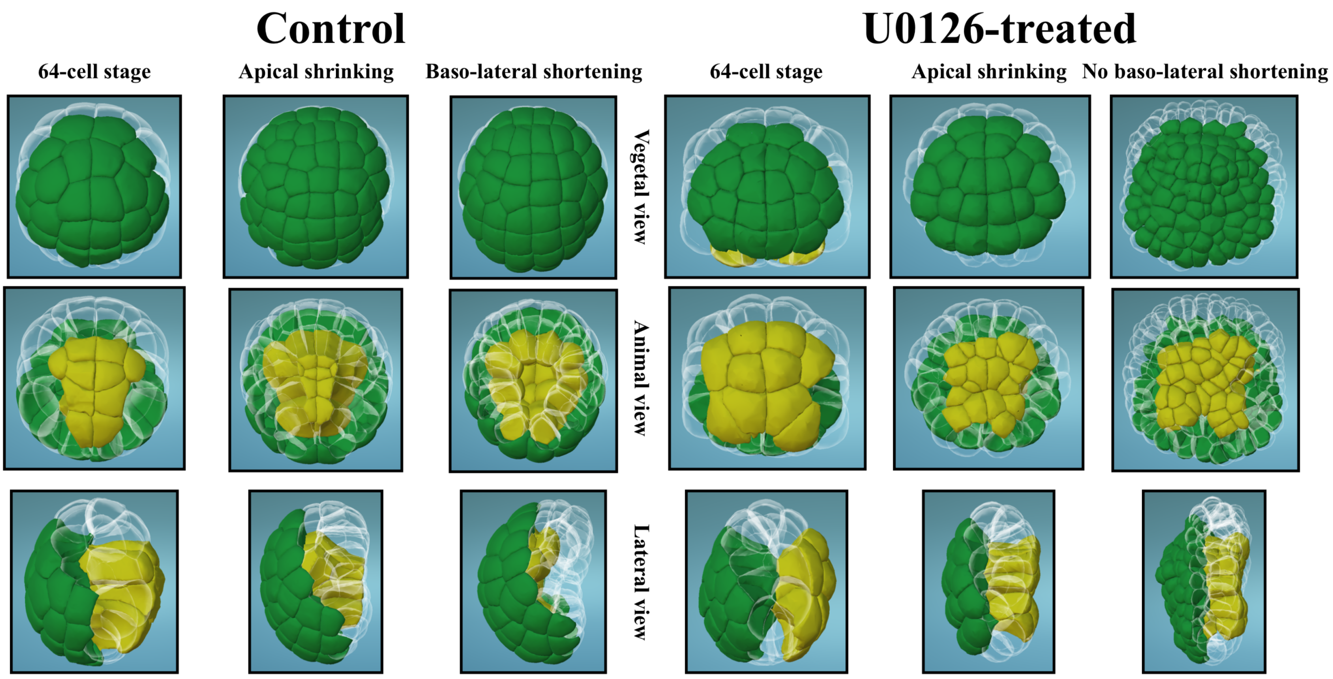


FIG. 2: Reconstruction of gastrulating embryos of *P. mammillata*, in both control conditions [1] and after U0126 treatment. Epidermal cells are labeled in green and endodermal cells (actively changing their shape) in yellow. Transparent cells belong to other embryonic tissues. Data visualized in MorphoNet [4].

### III. *P. MAMMILLATA* EMBRYONIC EPIDERMAL CELLS AND ANALYSIS OF THEIR DIVISION DIRECTIONS

In order to calculate main elongations and division directions of cells in the embryonic epidermis of *P. mammillata*, we started from the cell-level 3D volume reconstruction of two whole embryos produced in [1].

On these embryos, thanks to their invariant lineage, cells over several generations can be unambiguously named [2, 3]. At the 64-cell stage, symmetric pairs of cells a7.11, a7.12, a7.14, a7.15, a7.16, b7.11, b7.12, b7.13, b7.14, b7.15 and b7.16 were selected, corresponding to cells who will give rise to the embryonic epidermis after fate determination (green cells in Fig. 2).

Each of these cells and their epithelial progeny for up to three rounds of division have been selected along the timestack of 3D volume reconstruction of each of the two analysed embryos.

It being known that division of these cells always happens within the local tangential plane given by each cell's apical surface, to determine their main elongation we first selected their apical surfaces only out of the full 3D information on cell shapes.

The division orientation of each cell is defined as the direction joining the center-of-mass of the voxels constituting the apical surfaces of the two daughters of the dividing cell, measured two to four minutes after completion of cytokinesis.

Next we wished to correlate this direction to the main interphase elongation of the cell. To this end, we measured the shape of the apical surface of each cell between 23 and 25 minutes before the onset of cytokinesis (since two minutes is the time resolution of our datasets). Given the fact that the length of cell cycle in epidermis is between 1 and 2 hours, this is enough for the cells to have achieved their rest shape and far enough from cytokinesis to avoid biases due to the start of anaphase elongation.

For each cell $i$, we performed a principal component analysis on its apical surface to determine the direction of its main elongation $\hat{\mathbf{l}}^{(i)}_{\text{apical}}$ and the position of its center of mass $\mathbf{b}^{(i)}_{\mathbf{apical}}$. The division direction was then defined as the direction $\hat{\mathbf{d}}^{(i)}_{\text{apical}}$ of the line joining the apical centers of mass $\mathbf{b}^{(i_1)}_{\mathbf{apical}}$ and $\mathbf{b}^{(i_2)}_{\mathbf{apical}}$ of its two daughter cells $i_1, i_2$. The directions $\hat{\mathbf{l}}^{(i)}_{\text{apical}}$ and $\hat{\mathbf{d}}^{(i)}_{\text{apical}}$ are sufficient to calculate the parameters $\Delta_{\text{HR}}, \alpha$ and $\gamma$ shown in Figures 3 and 4 of the main text.

### IV. GEOMETRICAL PROPERTIES OF EPIDERMAL PRECURSORS IN DEVELOPING ASCIDIAN EMBRYOS: A QUASI-2D SYSTEM

#### A. Dynamics of 3D shapes of ascidian epidermal precursors

To prove that the morphogenesis of embryonic ascidian epidermis is a fit system to be explored by the simulated system of Fig. 3(a) of the main text, we have investigated whether the two main hypotheses of the model are valid there. The first hypothesis is that the process be, to a good degree of approximation, describable as a 2D process.To test this hypothesis, we have assessed how far from a planar shape the apical surface of each clone is. To this end, we have extracted, for every clone of every epidermal precursors at the 64-cell stage and for both embryos, the set of voxels composing their total apical surface during interphase. In this case, the interphase was defined slightly different than what done for single-cell shape analyses, since here we had the need to define an interphase period for the whole collection of cells in the clone. Hence, given a set $\{c\}$ of cells in a clone, we have defined they common cycle interval $T_c$ as the largest timespan at which all cells in $\{c\}$ exist (i.e., have been already produced by the division of their mothers and have not yet divided themselves). Clonal interphase was defined as the central timepoint of the interval $T_c$.

For each of the apical voxel sets, we then calculated the best-approximating plane, i.e., the plane minimizing the average distance of each voxels to it. We have then compared the average voxel distance to the plane $\mathcal{C}_{\text{clone}} = \frac{1}{N_{\text{voxels}}} \sum_{v \in \text{voxels}} d(v, \text{plane})$ to the main elongation of the apical clonal surface $\mathcal{E}_{\text{clone}}$, obtained through a principal component analysis of the voxel set: if the clone apical surface is non-negligibly curved with respect to its linear dimension, the average voxel distance to the plane must be non-negligible with respect to the main surface elongation. In other words, we assessed whether the curvature of the surface was comparable to the surface elongation through the parameter $\frac{\mathcal{C}_{\text{clone}}}{\mathcal{E}_{\text{clone}}}$, shown in Fig. 3 for each clone for both Control embryos. Values are averaged over all rounds of cell divisions in the clone. One easily sees that $\frac{\mathcal{C}_{\text{clone}}}{\mathcal{E}_{\text{clone}}}$ is less than 0.01 for all clones, i.e., the average distance of the surface from its planar approximation is less than 1/100 the elongation of the clone. At these scales, clones can be regarded as being approximately planar during the period of observation.

The second hypothesis is the fact that, during mitosis, cells significantly change their shape. This was investigated by looking at the change of the size of epidermal apical area. Of course, given the constraint of constant total volume, such increase in apical surface means a shortening in the baso-lateral direction. One can understand these changes through a simple toy model, represented in Fig. 4: assume a cell is a regular cuboid, whose apical surface is a square of edge $l_0$ and with a baso-lateral edge of length $k_0 l_0$. When dividing, the cell produces two regular cuboids, of apical edge $l_1$ and baso-lateral edge $k_1 l_1$. It is straightforward to see that, if the shape of the two daughters is the same as the shape of the mother modulo a global scaling factor (i.e., if $k_0 = k_1$), then $S_{\text{daughters}} = 2^{1/3} S_{\text{mother}}$, $S_{\text{daughters}}$ and $S_{\text{mother}}$ being, respectively, the total apical surfaces of the two daughters and the apical surface of the mother. More in general, this relation is

$$S_{\text{daughters}} = 2^{1/3} \left( \frac{k_0}{k_1} \right)^{2/3} S_{\text{mother}}, \tag{7}$$

which means that, if the ratio of apical to baso-lateral linear dimensions of daughters is lower (higher) that that of the mother, the scaling is faster (slower) than $2^{1/3}$. Fig.4 shows that this simple model accounts well for the observed changes in shape during divisions, represented by one typical example in the bottom part of the figure. The scaling roughly goes as $2^{1/2}$ for cells largely reducing their baso-lateral elongation during division (first division), to $2^{1/3}$ for cells changing their shape only by a global scaling factor (second division), to a value close to $2^{1/6}$ for the third division, in agreement with the shape change represented in the figure.

#### B. Dynamics of apical surfaces of ascidian epidermal precursors

Once established that ascidian epidermis can be regarded as a quasi-2D system, we analyse here the shape dynamics of the apical surface of ascidian epidermal precursors. Starting from segmented datasets of the two wild-type embryos we selected, for each epidermal cell starting from the 64-cell stage and until the end of each movie, the voxels composing the apical cellular surface. On each of these apical voxel sets we performed a principal component analysis, from which the ratio $r = \frac{\lambda_2}{\lambda_1}$ of two first principal values $\lambda_1 \geq \lambda_2$ was extracted. This ratio measures the elongation of the cell

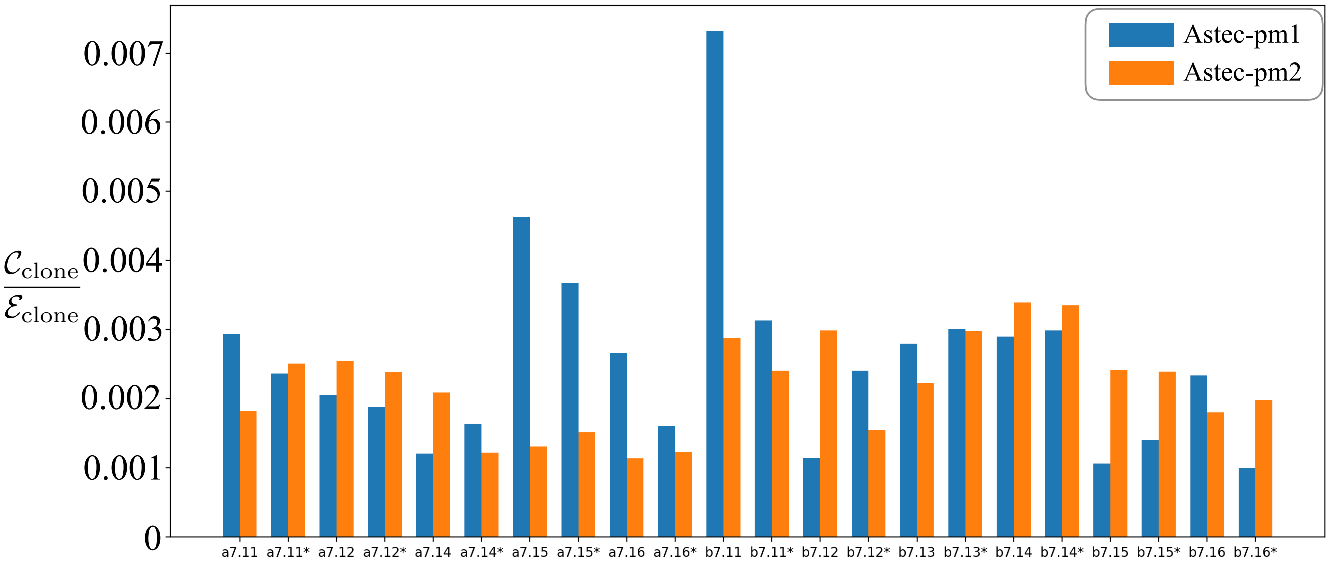


FIG. 3: Departure from planar geometry for each clone in both Control embryos, measured as the ratio $\frac{\mathcal{C}_{\text{clone}}}{\mathcal{E}_{\text{clone}}}$, where $\mathcal{C}_{\text{clone}}$ is the average distance of voxels composing the clonal apical surface surface from the plane that better approximates the surface; and $\mathcal{E}_{\text{clone}}$ is the main elongation of the clonal apical surface. Both values are time-averaged over the dynamics of clone shape following epidermal cell divisions.

in its apical place: $r \ll 1$ means that the apical surface of the cell is highly elongated in one direction, while $r = 1$ means that the apical surface has no preferential direction of elongation.

Since each cell cycle lasts over several timepoints, this procedure gave us dynamic information on the apical surface shape as a collection of ratios $\{r_c^{(k)}\}$, with $c$ labelling each cell and $k$ characterizing the specific timepoint at which these values have been measured.

In order to compare cells with different cell-cycle length, we replaced the timepoint index by a percentage measure of the progression of the cell cycle, 0% being the moment the cell is born and 100% being the moment its cytokinesis is completed.

By linear interpolation between two values at two consecutive timepoints, we could hence characterise the continuous functions $r_c(p)$, where $c$ labels the specific cell and $p \in [0, 100]$ is the percentage of its cell cycle.

In addition we considered the parameter

$$g_c(p) = \frac{r_c(p) - r_c(100)}{r_c(100)}, \tag{8}$$

quantifying the shape dynamics as a variation relative to the final shape of the cell just at the end of its cytokinesis. This normalisation allows us to better compare the dynamics of cells having different values of their elongation.

Finally, we define the two parameters

$$\Xi(p) = \langle r_c(p) \rangle_c\,, \tag{9}$$
$$\Gamma(p) = \langle g_c(p) \rangle_c\,, \tag{10}$$

as the average values over all epidermal cells in the two datasets (for which a full cycle information is available) of the elongation ratio $r_c(p)$ and its relative variation $g_c(p)$.

These parameters as a function of $p$ are shown, respectively, in Fig. 5 and Fig. 6, together with their respective standard deviations (grey area). While the first has a direct geometrical meaning (the higher $\Xi$, the lower the elongation on average), the second removes some artefacts of the averaging procedure due to the broad distribution of shape elongations.

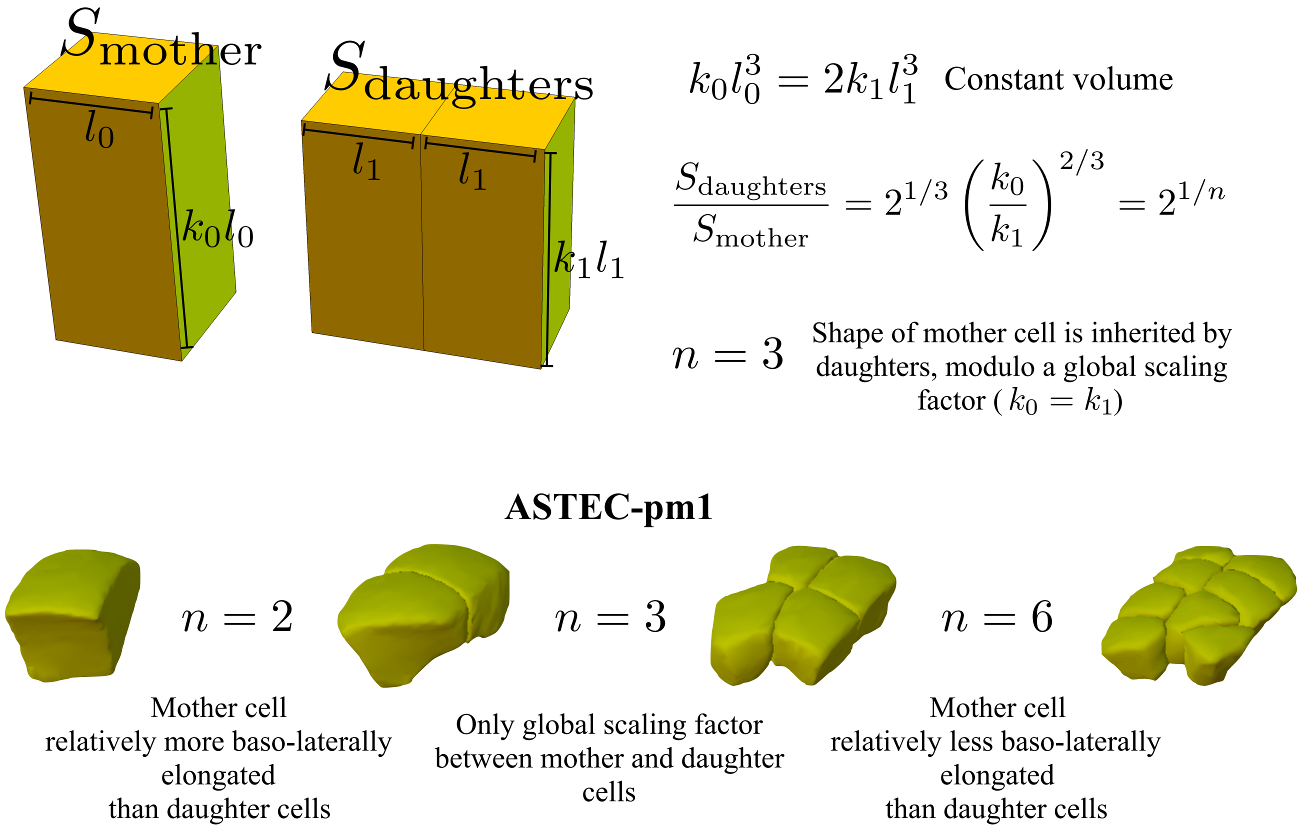


FIG. 4: Schematic view of apical surface changes after cell division. Cuboid cells produce cuboid cells of constant total volume. This produces apical surface growth roughly by a factor $2^{1/n}$: as shown in (7), $n$ depends on the change in ratio between the baso-lateral extension of the cell and its apical elongation. For instance, $n = 3$ if the basolateral to apical ratio of the mother is perfectly conserved in the daughters, $n < 3$ if mothers cells are more elongated in the baso-lateral direction than daughter cells and $n > 3$ in the opposite case. This is close to what is observed, as illustrated by a typical shape dynamics in ASTEC-pm1. See that for instance the shape of cells before and after the second division round does not change much, apart for a global scaling factor: this is well recapitulated by the fit at $n = 3$.

In both cases, one notices an increase of the parameter towards the end of the cycle, corresponding to the apical mitotic rounding (green line in the insets), followed by a sharp decrease as the cell progresses through anaphase and cytokinesis (red line in the insets). It is evident that mitotic rounding entails a smaller shape change than the subsequent anaphase elongation. In addition, the shape dynamics linked to mitotic rounding is slower that the following elongation, since it produces smaller shape changes distributed over a larger portion of the cell cycle. Hence, the energy dissipated during mitotic rounding due to viscosity of the cellular medium is arguably smaller than the energy dissipated during the elongating phase.

Also, from Figures 5 and 6 one notices an initial increase in both $\Xi$ and $\Gamma$ in the very first phases of the cell cycle (low progression percentages). This corresponds to the (roughly) round shape attained by the cell at the end of the cytokinesis of its mother cell. This initial increase of $\Xi$ and $\Gamma$ is due to the final part of the trajectory followed by the mother cell during division, and justifies our assumption in Section II of regular polygonal shape of daughter cells.

#### C. Volume ratio of daughter cells of ascidian epidermal precursors

We analyse here the distribution of the ratios of the volumes of each pair of epidermal sister cells for the two datasets of WT ascidian development. Figure 7 shows the distribution of the ratio

$$R_v = \frac{V_{\min}}{V_{\max}}, \tag{11}$$

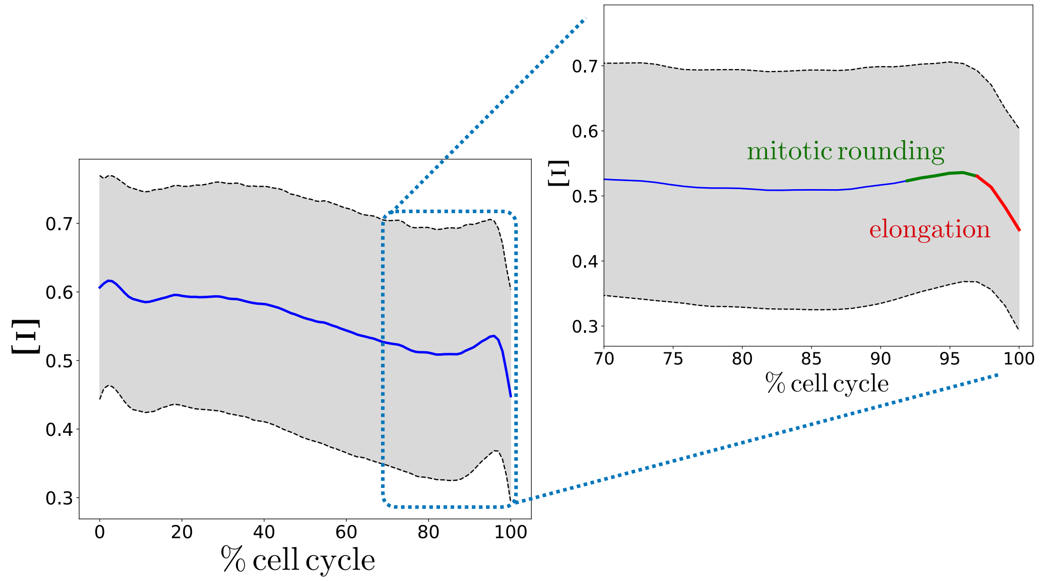


FIG. 5: Average elongation ratio Ξ as a function of the percentage of cell-cycle progression (blue line), together with its standard deviation (grey area). In the inset, showing a magnification around the end of the cycle, the increase due to mitotic rounding and the decrease due to anaphase and cytokinesis elongation are highlighted in, respectively, green and red.

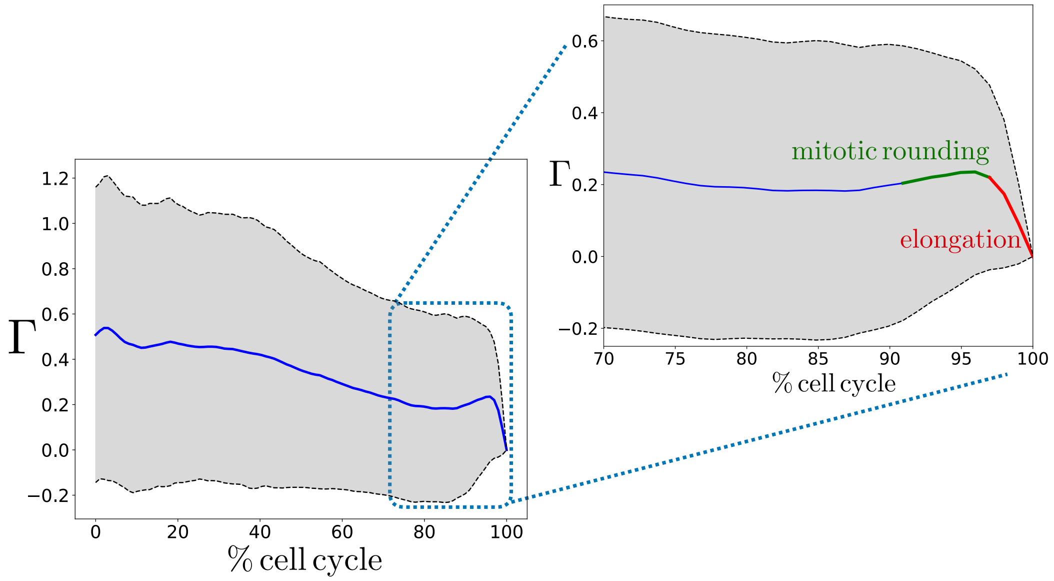


FIG. 6: Average relative variation of the elongation ratio Γ as a function of the percentage of cell-cycle progression (blue line), together with its standard deviation (grey area). In the inset, showing a magnification around the end of the cycle, the increase due to mitotic rounding and the decrease due to anaphase and cytokinesis elongation are highlighted in, respectively, green and red.

where $V_{\min}$ and $V_{\max}$ are, respectively, the volume of the smaller and of the bigger of the two sister cells. The distribution is obtained from the analysis of 638 divisions in the epidermis of the two ascidian datasets ASTEC-pm1 and ASTEC-pm2.

One sees that the great majority of divisions in ascidian epidermis produces daughter cells of similar volumes (59%

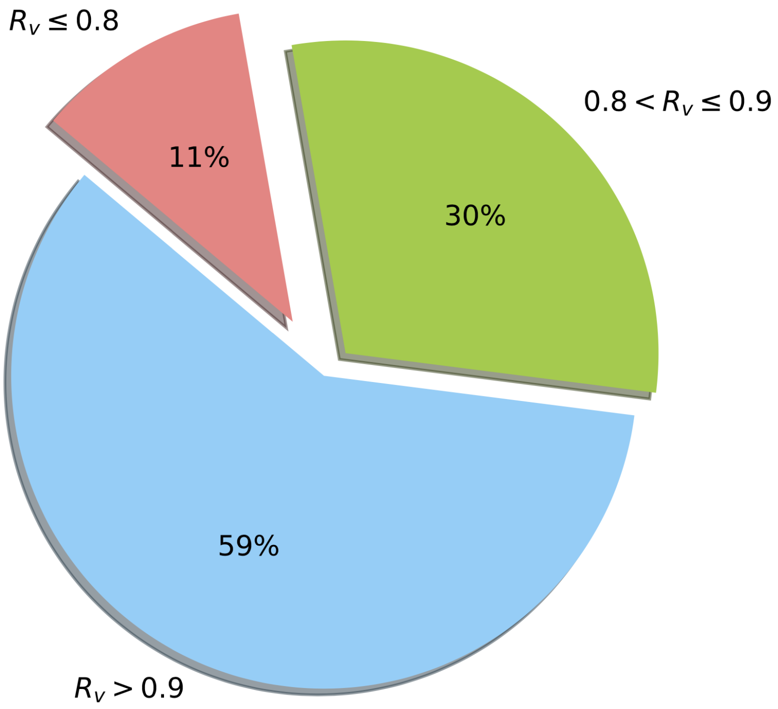


FIG. 7: Distribution of sister cells volume ratio $R_v$ defined in Eq. (11), for 638 cell divisions in ascidian epidermis of the two WT datasets ASTEC-pm1 and ASTEC-pm2.

of divisions have $R_v > 0.9$, and 89% of divisions have $R_v > 0.8$). Since the total embryonic volume does not change, two sister cells of equal volume can only be produced by geometrically-symmetric divisions of their mother, which justifies our assumption in Section II according to which the division plane passes through the geometric center of the dividing cell.

### V. U0126-TREATED *P. MAMMILLATA* EMBRYO AND ITS PHENOTYPE

Inhibition of the FGF/ERK signalling was previously shown to impair gastrulation [5] by a fate switch of A-line endoderm precursors to trunk lateral cells [6] that impair and block the invagination during gastrulation as shown in Fig. 2 For FGF/ERK signalling inhibition, Phallusia mammillata embryos were treated with U0126 (MEK inhibitor; abcam:ab120241) at a concentrations of 2uM diluted in artificial sea water [5], from the early 16-cell stage.

[1] L. Guignard, U.-M. Fiuza, *et al.*, *Contact-dependent cell-cell communications drive morphological invariance during ascidian embryogenesis*, bioRxiv: https://doi.org/10.1101/238741 (2017).
[2] E. G. Conklin, *The organization and cell - lineage of the ascidian egg* (Philadelphia :[Academy of Natural Sciences], 1905).
[3] M. Brozovic *et al.*, *ANISEED 2017: extending the integrated ascidian database to the exploration and evolutionary comparison of genome-scale datasets.*, Nucleic Acids Res. **46**, D718 (2018).
[4] B. Leggio, J. Laussu, A. Carlier, C. Godin, P. Lemaire, and E. Faure, *MorphoNet: an interactive online morphological browser to explore complex multi-scale data*, Nature Comm. **10**, 2812 (2019).
[5] C. Hudson *et al.*, *A conserved role for the MEK signalling pathway in neural tissue specification and posteriorisation in the invertebrate chordate, the ascidian Ciona intestinalis*, Dev. **130**, 147 (2003).
[6] W. Shi and M. Levine, *Ephrin signaling establishes asymmetric cell fates in an endomesoderm lineage of the Ciona embryo*, Dev. **135**, 931 (2008).

TABLE I: Parameters and variables of the model

| Parameter | Formula | Description |
|---|---|---|
| $\eta$ | - | Viscous friction coefficient of vertex displacement in the cellular medium |
| $\hat{\mathbf{n}}$ | - | Division orientation (direction perpendicular to division plane) |
| $\mathbf{v}_k^{(i)}$ | - | Velocity vector of the $i$-th vertex of daughter $k$ |
| $\mathbf{p}_k^{(i)}$ | - | Position vector of the $i$-th vertex of daughter $k$ |
| $\tau_k^i$ | - | Trajectory followed by the $i$-th vertex of daughter $k$ |
| $\mathbf{F}_\sigma$ | $\psi_\sigma(\mathbf{r}, \mathbf{r}_\sigma)\hat{\mathbf{w}}(\mathbf{r}, \mathbf{r}_\sigma)$ | Force vector exerted at position $\mathbf{r}$ by an active process $\sigma$ at position $\mathbf{r}_\sigma$ |
| $\hat{\mathbf{w}}(\mathbf{r}, \mathbf{r}_\sigma)$ | - | Force direction at position $\mathbf{r}$ by the active process $\sigma$ |
| $\psi_\sigma(\mathbf{r}, \mathbf{r}_\sigma)$ | $F_\sigma h\big(d(\mathbf{r}, \mathbf{r}_\sigma)\big)$ | Force intensity at position $\mathbf{r}$ by the active process $\sigma$ |
| $h\big(d(\mathbf{r}, \mathbf{r}_\sigma)\big)$ | Monotonically decreasing function | Function describing the decrease of force magnitude with the distance $d$ |
| $\phi_\sigma$ | $\frac{F_\sigma}{\eta}$ | Ratio of maximal force intensity to Viscous friction coefficient |
| $\lambda_c$ | - | Interphase cellular cortical tension |
| $\kappa_c$ | - | Interphase cellular surface elasticity |
| $\rho$ | $\frac{\phi_\sigma}{v}$ | Ratio of active to viscous forces in case of constant velocity $v$ |
| $\zeta$ | $\frac{L_d}{L_0}$ | Ratio of stress penetration length to linear size of multicellular system |
| $\alpha$ | $1 - \langle \chi_{\mathrm{CJ}} \rangle / 90$ | Average deviation from geometrical rule for division orientation |
| $\gamma$ | $\big\langle \lvert \mathbf{n}_t^{(c)} \cdot \mathbf{n}_{t+1}^{(c)} \rvert \big\rangle$ | Average correlation between consecutive division orientations |
